# Epidemiology and phylodynamic analysis of canine distemper virus circulating in Michigan, USA

**DOI:** 10.64898/2026.06.03.729155

**Authors:** Kota Nakasato, Julie Melotti, Scott Fitzgerald, Lindsy Hengesbach, Steffanie Anderson, Kayla Conner-Halim, Annabel Wise, Danielle Thompson, Tuddow Thaiwong-Nebelung, Kimberly Dodd, Roger Maes, Scott Sherrill-Mix

**Affiliations:** Veterinary Diagnostic Laboratory, Michigan State University, MI; Wildlife Disease Laboratory, Michigan Department of Natural Resources, MI; Department of Microbiology, Genetics, & Immunology, Michigan State University, MI; Interdepartmental Microbiology Graduate Program, Iowa State University; Department of Plant Pathology, Entomology, and Microbiology, Iowa State University

**Keywords:** CDV, canine distemper virus, Nextstrain, Michigan, wild mesocarnivores, domestic dogs

## Abstract

Canine distemper virus (CDV) is a highly contagious generalist pathogen that can cause significant mortality in domestic dogs and wild mammals. Due to the fast-evolving nature of CDV and its global spread, epizootic events and the emergence of new strains have been frequently reported. In Michigan, CDV surveillance from 2008 to 2018 was previously reported. Here, we combine and extend these data through 2023 to bring together 16 years of CDV surveillance in wild mammals in Michigan. We also sequenced CDV strains originating from both wildlife and domestic dogs to examine viral evolution across host populations. To facilitate interpretation of these data in both local and global contexts, we developed a Nextstrain workflow for CDV, enabling interactive visualization of viral evolution over time and geographic space. Our data show persistence of CDV in Michigan mammals during the study period and point to temporal, geographic, and host factors associated with CDV occurrence. Phylogenetic analysis using the newly built Nextstrain workflow showed that three CDV lineages—America-3, America-5, and Canada-1—are currently circulating in both wild and domestic animals in Michigan. The Nextstrain workflow enables reproducible, scalable integration of genomic sequencing into local surveillance and provides an updateable platform for ongoing and future surveillance efforts. This study demonstrates the value of coupling wildlife surveillance and diagnostic testing with genomic sequencing to identify lineage turnover and anticipate changes in viral behavior.

## Introduction

Canine distemper virus (CDV; *Morbillivirus canis*) is a highly contagious pathogen that causes significant morbidity and mortality across a broad range of carnivore species, including *Canidae, Mustelidae*, and *Procyonidae* [1]. CDV is a member of the family *Paramyxoviridae* in the genus *Morbillivirus* and is related to other important pathogens causing public health and economic concerns, such as measles and rinderpest [2]. CDV is the etiological agent of canine distemper in domestic dogs. Clinical signs of canine distemper include ocular and nasal discharge, respiratory distress, diarrhea, vomiting, and neurological impairment that can be confused with those of rabies [3]. The ability of CDV to infect lymphoid, epithelial and central nervous system cells underlies its broad tropism and diverse clinical manifestations [4]. A safe and efficacious vaccine is available for domestic dogs, and vaccination is strongly recommended by veterinary authorities [5].

As a pathogen infecting more than 100 species of mammals belonging to 12 families, CDV also poses a substantial threat to many species of wildlife globally [1, 6]. In Tanzania’s Serengeti National Park, for example, over 85% of lions were infected in a single outbreak event in 1994, leading to approximately 30% mortality in the population [7]. CDV is also a risk for endangered felid species such as Amur leopards (*Panthera pardus orientalis*), Amur tigers (*Panthera tigris altaica*), and Javan leopards (*Panthera pardus melas*) [8–10]. In marine species, mass mortality among seals populations in Lake Baikal and the Caspian Sea has been attributed to CDV [11]. Although not known to infect humans, some strains of CDV have been reported to infect non-human primates and cause outbreaks with significant mortality in rhesus monkeys (*Macaca mulatta*) in China and cynomolgus monkeys (*Macaca fascicularis*) in Japan [12, 13].These cases underscore CDV’s capacity to cross species barriers and threaten the conservation of species already at risk.

CDV has a linear single strand negative-sense RNA genome which is approximately 15.7kb long encoding several structural and nonstructural proteins [4]. Playing a central role in host specificity and viral evolution, the hemagglutinin (H) protein mediates viral attachment by binding to host receptors such as SLAM and Nectin-4 and is under strong selective pressure by the host immune system and receptor variation [4]. As a result, the H gene exhibits the highest genetic variability within the CDV genome and serves as the primary marker for phylogenetic classification [14]. Based on phylogenetic studies of H gene sequence, CDV are generally grouped into geographically distinct lineages [15–20]. Currently available vaccines are mostly derived from the America-1 lineage and have been in widespread use since the 1950s and 1960s [21]. Although recent structural analysis of CDV H provides a possible explanation for the continued effectiveness of older vaccine strains [22], some cases of non-America-1 lineages causing clinical signs in vaccinated dogs have been reported [23, 24].

The rapid evolution and antigenic variability of CDV highlight the importance of genomic surveillance to detect existing and emerging CDV variants and understand their movement across species and geographic boundaries. Despite increasing availability of CDV genomic data, there is currently no standardized framework for integrating and visualizing these data. Advances in sequencing technologies and open-source analytical platforms, such as Nextstrain, now make it feasible to track the evolution of human and animal pathogens dynamically [25]. Applying such approaches to CDV will improve our ability to monitor lineage emergence, cross-species transmission, and regional spread, ultimately strengthening wildlife disease management and guiding new vaccination strategies.

In Michigan, the Wildlife Disease Laboratory (WDL) of the Michigan Department of Natural Resources (MI DNR) has monitored CDV in wild carnivores showing clinical signs of potential infection since 2008 [26]. Fitzgerald et al. [26] summarized surveillance data from 2008–2018 and reported a marked increase in CDV-positive wild carnivores detected in the mid-2010s. Sequencing of three viral isolates revealed the circulation of a lineage previously labeled “America-2” along with a unique Michigan-specific sequence type. Building on this foundation, we extended CDV surveillance of wild mammals in Michigan through 2023 and integrated molecular data from both wildlife and domestic dogs to explore the virus’s epidemiology in local and global contexts. Specifically, we investigated: (1) temporal and geographic trends in CDV detection in affected wild species across Michigan; (2) genetic relationships between viral sequences from wild and domestic hosts to assess evidence of cross-species transmission; and (3) the emergence and evolutionary dynamics of novel or introduced CDV variants circulating in the region. To facilitate continued monitoring and data sharing, we developed an interactive Nextstrain workflow for CDV, providing an open platform for real-time visualization and phylodynamic tracking of this globally important pathogen.

## Methods

### Wild carnivore samples and data collection

Sample and data collection methods followed procedures previously described by Fitzgerald et al. [26]. Briefly, wild mammals exhibiting respiratory, enteric, or neurological signs were reported by the public and collected by personnel from the MI DNR. Animals found alive were humanely euthanized. Carcasses were frozen prior to transport to the WDL. At the WDL, species, sex, approximate age, collection and/or receipt date and county of origin were recorded. A full postmortem examination was conducted, during which appropriate tissues were collected, fixed in 10% neutral buffered formalin and paraffin embedded. Fresh tissue samples from some individuals were stored at −20 °C for RT-PCR and sequencing.

For each submitted case, high priority zoonotic pathogens, including rabies virus and H5N1 highly pathogenic avian influenza when suspected, were tested first. Rabies testing was performed using the direct rapid immunohistochemical test (DRIT) by the USDA Wildlife Services. Testing for H5N1 was conducted by the Michigan State University Veterinary Diagnostic Laboratory (MSU VDL). Following exclusion of these pathogens, tissue sections were examined histologically and by immunohistochemistry (IHC) [27].

New data from 2019-2023 was combined with the data from 2008-2018 reported by Fitzgerald et al. [26]. Most submissions were tested for CDV by IHC (Table 1). Untested cases arose due to intermittent funding constraints or strong evidence of alternative etiologies. For these untested submissions, cases with necropsy and histopathological findings consistent with CDV (e.g., characteristic lesions in brain or lung) were classified as probable positives. Untested cases lacking CDV-compatible lesions or those with compelling evidence of another cause of illness (e.g., other pathogens or toxins identified) were classified as putative negatives. To assess the impact of uncertainty in case classification, all analyses were performed: (i) using the full dataset, including probable positives and putative negatives, and (ii) using only IHC-tested cases.

**Table 1.**
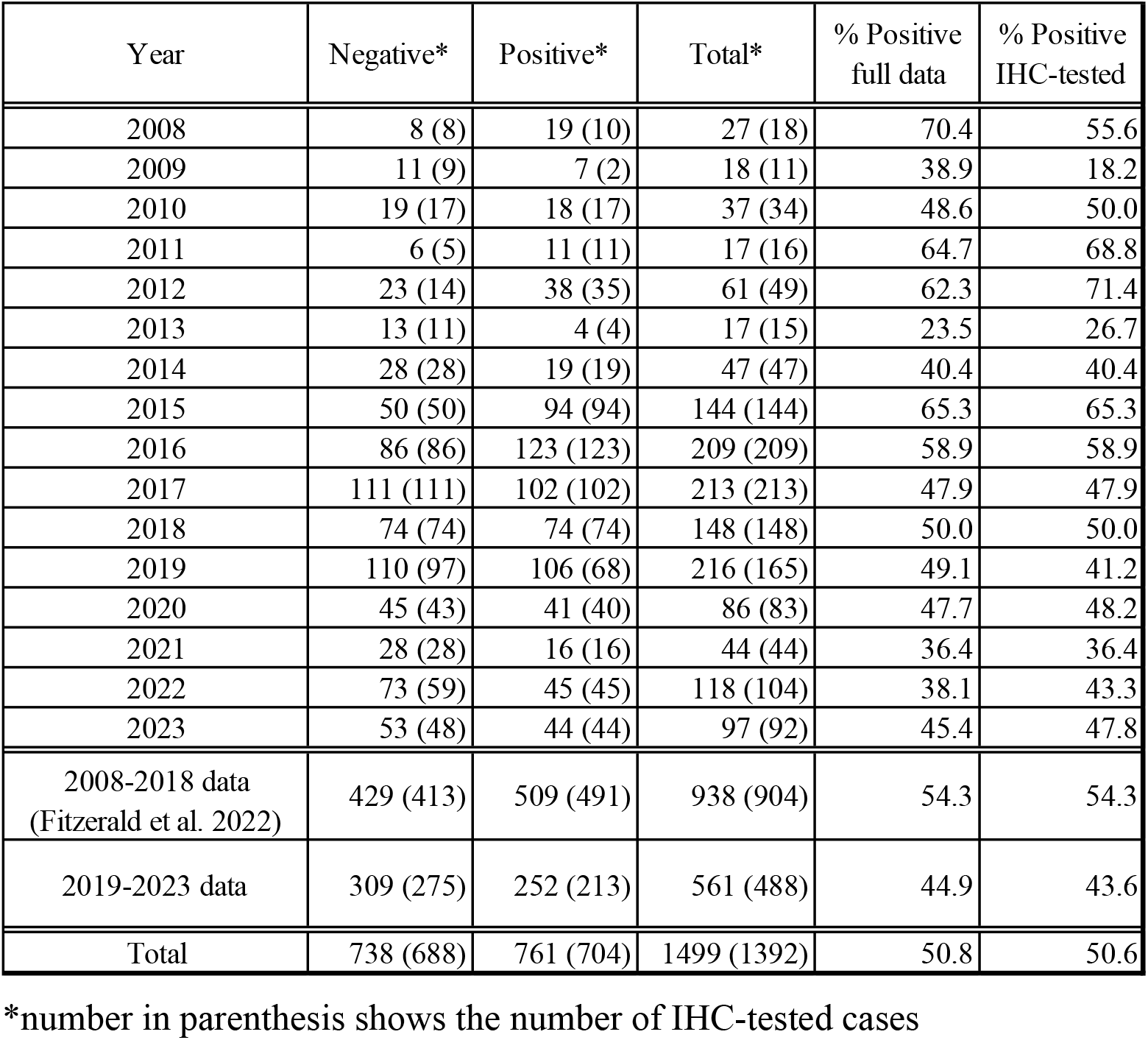
Summary of CVD case submissions and test results in Michigan from 2008 to 2023.

| Year | Negative* | Positive* | Total* | % Positive full data | % Positive IHC-tested |
| --- | --- | --- | --- | --- | --- |
| 2008 | 8 (8) | 19 (10) | 27 (18) | 70.4 | 55.6 |
| 2009 | 11 (9) | 7 (2) | 18 (11) | 38.9 | 18.2 |
| 2010 | 19 (17) | 18 (17) | 37 (34) | 48.6 | 50.0 |
| 2011 | 6 (5) | 11 (11) | 17 (16) | 64.7 | 68.8 |
| 2012 | 23 (14) | 38 (35) | 61 (49) | 62.3 | 71.4 |
| 2013 | 13 (11) | 4 (4) | 17 (15) | 23.5 | 26.7 |
| 2014 | 28 (28) | 19 (19) | 47 (47) | 40.4 | 40.4 |
| 2015 | 50 (50) | 94 (94) | 144 (144) | 65.3 | 65.3 |
| 2016 | 86 (86) | 123 (123) | 209 (209) | 58.9 | 58.9 |
| 2017 | 111 (111) | 102 (102) | 213 (213) | 47.9 | 47.9 |
| 2018 | 74 (74) | 74 (74) | 148 (148) | 50.0 | 50.0 |
| 2019 | 110 (97) | 106 (68) | 216 (165) | 49.1 | 41.2 |
| 2020 | 45 (43) | 41 (40) | 86 (83) | 47.7 | 48.2 |
| 2021 | 28 (28) | 16 (16) | 44 (44) | 36.4 | 36.4 |
| 2022 | 73 (59) | 45 (45) | 118 (104) | 38.1 | 43.3 |
| 2023 | 53 (48) | 44 (44) | 97 (92) | 45.4 | 47.8 |
| 2008-2018 data<br>(Fitzerald et al. 2022) | 429 (413) | 509 (491) | 938 (904) | 54.3 | 54.3 |
| 2019-2023 data | 309 (275) | 252 (213) | 561 (488) | 44.9 | 43.6 |
| Total | 738 (688) | 761 (704) | 1499 (1392) | 50.8 | 50.6 |

\*number in parenthesis shows the number of IHC-tested cases

### RNA extraction, qRT-PCR, and Sanger Sequencing

A subset of samples collected in 2022 and 2023 were used for RNA extraction, RT-PCR, and sequencing of the hemagglutinin gene for phylogenetic and genotype analysis. RNA was extracted from fresh frozen tissues using the Qiagen RNA Extraction Kit (Qiagen, Germany), and probe-based reverse transcription Taqman-probe quantitative PCR (qRT-PCR) was performed to confirm CDV-positive cases at the Virology Section of the MSU VDL [28]. CDV H gene sequences were amplified and sequenced using a combination of published and newly designed primer sets (Table S1). RT-PCR was conducted using the Qiagen One-Step RT-PCR Kit (Qiagen, Germany) or a combination of High-Capacity cDNA Reverse Transcription Kit (ThermoFisher Scientific, USA) with Taq DNA Polymerase (Qiagen, Germany). The resulting PCR products were gel-purified using QIAquick Gel Extraction Kit (Qiagen, Germany). Purified products were sequenced either at the Next Generation Diagnostic Section of MSU VDL with a SeqStudio Genetic Analyzer (ThermoFisher Scientific, USA) or submitted to the Research Technology Support Facility Genomics Core at MSU.To investigate the circulation of CDV between wild species and domestic dogs in Michigan, selected domestic dog samples submitted to the MSU VDL Virology sections for CDV testing were also sequenced. In addition, three samples from the prior study [26] were also re-sequenced to obtain full length H gene sequences.

### Sequencing with Oxford Nanopore Technologies MinION

Four wild mesocarnivore samples were processed for metagenomic sequencing using the Oxford Nanopore Technology (ONT) MinION platform to generate longer genomic regions of CDV. Sequence-independent single-primer amplification (SISPA) was performed as previously described [29]. Briefly, viral nucleic acids were reverse-transcribed using tagged random primers, converted to double-stranded cDNA, and amplified by PCR prior to library preparation with the ONT Native Barcoding Kit. Sequencing was performed on R10.4.1 flow cells using a MinION device at at MSU VDL. Base-calling was performed with MinKNOW (version 23.07.15) and Guppy (version 7.1.4), and the resulting FASTQ files were analyzed on High Performance Computing Center resources offered by Institute for Cyber-Enabled Research at MSU. Read quality was assessed using NanoPlot [30] and NanoQC [31]. Chopper [30] was used to trim low-quality bases and adapter sequences, and Minimap2 [32] and SAMtools [33] were used to extract reads mapped to the reference CDV genome (NC_001921.1). These mapped reads were used for genome assembly. Due to the nature of the SISPA protocol, most reads were shorter than typical ONT sequencing reads (average read length across the four libraries: ∼1000 bp), which posed challenges for tools optimized for long-read data. Therefore, SPAdes with single-end read input was used for de-novo assembly [34]. The resulting contigs were compared with Sanger-sequenced fragments of the H gene (∼1000 bp), and base-level consistency between the ONT-generated contigs and Sanger sequences was confirmed. The assembled contigs were used for subsequent analyses.

### Development of CDV NextStrain workflow and phylodynamic analysis

For interactive and reproducible phylodynamic analysis, we built a Nextstrain workflow for CDV. All publicly available sequences and metadata annotated as “*Morbillivirus canis* (taxonomy id: 3052342)” were retrieved from NCBI Virus in November 2024. Within the workflow, sequences were aligned and H-gene datasets were generated by trimming the alignment to reference coordinates and retaining only sequences with adequate coverage across the targeted window. We first focused on building a dataset with full-length H gene sequences and used this dataset to optimize workflow parameters. In parallel, we created a workflow optimized for analyzing partial H gene sequences (∼900 bp; genomic positions 502–1401 based on the NC_001921.1 H-gene coordinates). Metadata fields relevant to downstream analyses (sampling date, host, and lineage/clade assignment) were curated from associated publications when available, and curation was conducted iteratively during workflow development to resolve missing or inconsistent records identified during preliminary phylogenetic reconstructions. In particular, sampling dates were updated based on previous phylodynamic studies [18, 20]. For records lacking a sampling date, the NCBI release date was used as an approximate estimate.

Aligned sequences used for phylogenetic tree construction were analyzed with RDP5 to detect potential recombinant strains [35]. The methods employed included RDP [36], GENECONV [37], BootScan [38], MaxChi[39], Chimaera [40], SiScan [41], and 3Seq [42]. A sequence was considered a potential recombinant if a recombination signal was detected by more than three of these methods and no warning flags were raised for false-positive by RDP5. Sequences from Michigan were not detected as recombinant sequences by this threshold.

After optimizing the workflow, CDV sequences from Michigan and their associated metadata were added to the dataset and analyzed to produce four phylogenetic trees: (1) full length H gene, (2) full length H gene without potential recombinant sequences, (3) partial H gene, and (4) partial H gene without potential recombinant sequences. To facilitate interpretation, selected clades of interest were visualized as zoomed-in subtrees using ggtree [43] for manuscript preparation.

### Statistical analysis of surveillance data

Data were visualized and analyzed using R (version 4.4.2) and RStudio (version 2025.09.0+387) [44, 45]. Descriptive statistics and figures were generated using tidyverse packages [46].

To investigate temporal, geographical and host-associated factors associated with CDV positivity, differences in proportions between groups were evaluated using a chi-squared test or Fisher’s exact test. Generalized linear models (GLMs) with a binomial distribution and logit link function were also used to assess the effects of biological, temporal, and spatial predictors on the likelihood of a positive test result. A linear regression model was used to evaluate factors associated with qRT-PCR Ct values.

For geographical analysis, we estimated the proportion of positive samples for each county using a Bayesian conditional autoregressive (CAR) model adapted from Marques et al. [47]. The model accounts for variable sampling effort across counties and spatial correlation between neighboring counties when estimating positivity.

## Results

### Surveillance of CDV in Michigan wildlife from 2008 to 2023

From 2019-2023, the MI DNR received 561 animals exhibiting abnormal behavior or signs of systemic illness suspected to be CDV. Combined with previously reported 2008-2018 data [26], the total dataset contains 1499 submissions from 82 counties and 17 wild animal species (Table 1). Overall, 92.9% (1392/1499) of the submissions in the dataset were tested by IHC testing; 7.1% of samples (n = 107) were not tested, but classified to probable positive or negative based on available evidence as described in the method section. To account for uncertainty, analysis were performed using both the full dataset (including putative positive and negative cases) and a subset limited to IHC-tested cases. The results were largely consistent between the two datasets; therefore, analyses based on the full dataset are presented below unless otherwise specified. The results from dataset limited to IHC-tested cases are available in supplementary data (Fig S1-2, 4).

The full dataset enabled a 16-year assessment (2008–2023) of CDV submissions and positivity in suspected wild species in Michigan. Over the study period, 704 samples tested positive for CDV and 57 were classified as probable positives. This corresponds to a positivity rate of 50.8% (761/1499) among suspect cases. As reported in Fitzgerald et al. [26], submissions and positive cases began to rise in 2016 and remained high through 2018. Submissions and positive cases were similarly elevated in 2019 but declined thereafter and remained below peak levels through 2023 (Fig 1a). The new dataset from 2019-2023 has a 44.9% (252/561) overall positivity rate, which decreased from the overall positivity rate of 54.3% (509/938) between 2008-2018 reported in Fitzgerald et al. [26] (Table 1).

**Figure 1.**
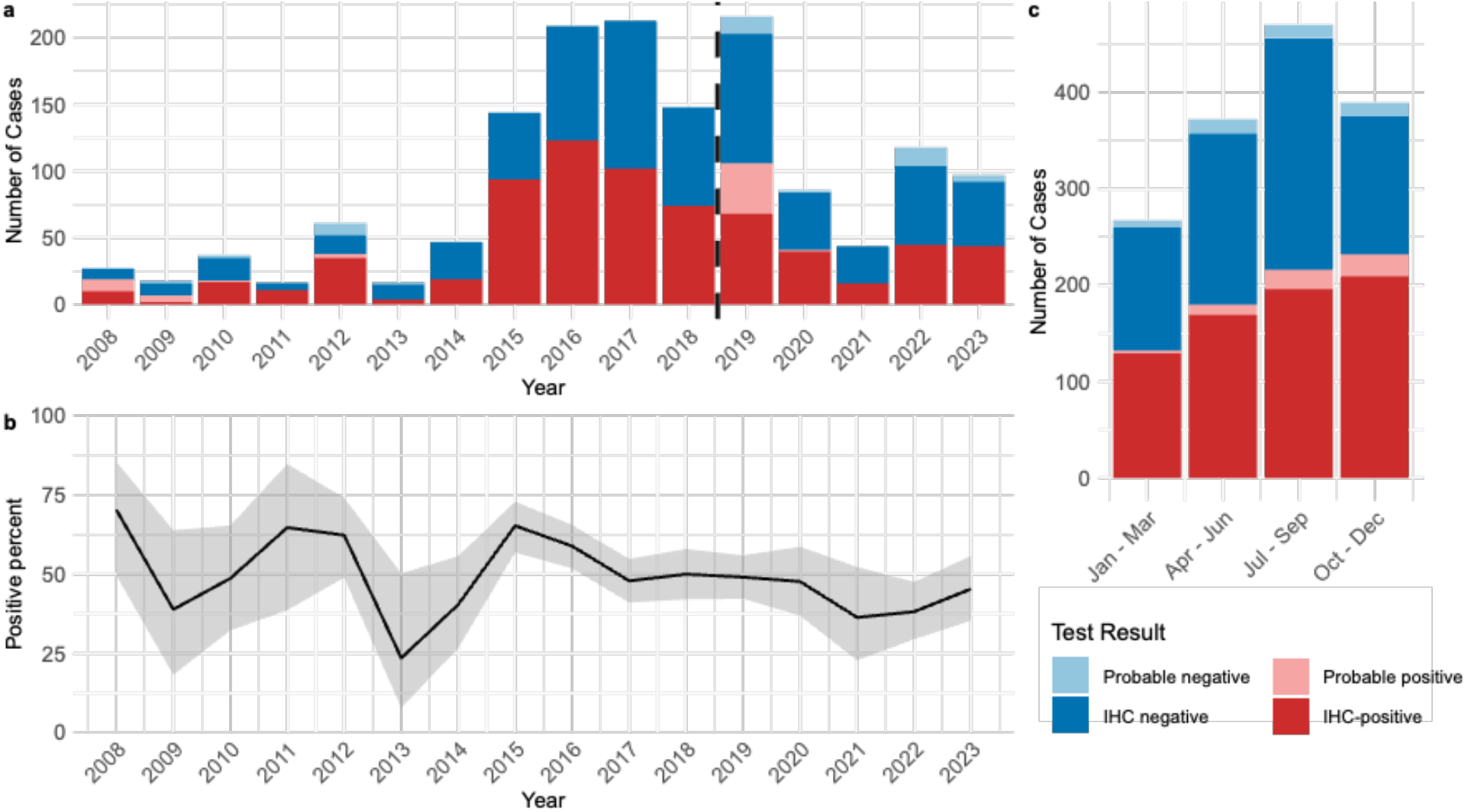
Temporal dynamics of CDV surveillance data from 2008 to 2023. (a) Annual numbers of wildlife submissions and immunohistochemistry (IHC) test results, showing counts of IHC-positive, IHC-negative and probable positive/negative cases. Dashed line indicates the break between cases previously reported in Fitzerald et al. [26] and cases newly reported here. (b) Temporal changes in CDV positivity, expressed as the proportion of positive and probable positives submissions among all tested samples with 95% confidence interval (gray shading). (c) Total number of submissions summarized by calendar quarter across the study period.

Annual CDV positivity rates varied across years (Fig 1b; Table 1), ranging from 23.5% in 2013 (95% CI: 7.8–50.2%; n = 17) to 70.4% in 2008 (95% CI: 49.7–85.5%; n = 27). Overlapping confidence intervals indicate that most early fluctuations were not statistically meaningful and likely reflect variability due to low submission numbers. After 2016, positivity rates stabilized around 40–50%, coinciding with larger and more consistent annual sample sizes. Overall, no pronounced directional trend was apparent over the 2008–2023 period. The quarterly season is significantly associated with positivity rates (X^2^ = 16.8, df =3, p = 0.0008). A pairwise comparison of proportions showed that October - December had a significantly higher positivity rate (59.6%) compared to April - June (48.4%; p = 0.04) and July-September (46%; p = 0.005) (Fig 1c).

Among the 17 wildlife species submitted, raccoons accounted for the majority of cases (60.5%, n = 907), followed by striped skunks (12.9%, n = 193), gray foxes (10.1%, n = 151), red foxes (8.0%, n = 120), coyotes (4.8%, n = 72), American mink (1.2%, n = 18), and gray wolves (1.0%, n = 15), with ten additional species represented by only a few submissions (Table 2). Thirteen species tested positive for CDV in at least one case. Positivity rates differed significantly among species (Fig 2a; Fisher’s exact test, p < 0.001). Among species with ≥10 submissions, gray foxes showed the highest positivity rate (78.8%, 95% CI: 71.3–84.9%), whereas red foxes had the lowest (7.5%, 95% CI: 3.7–14.2%). Raccoons had an intermediate positivity rate (56.2%, 95% CI: 52.9–59.5%), while striped skunks exhibited slightly lower rates (37.8%, 95% CI: 31.0–45.1%).

**Table 2.** Summary of CVD case submissions and test results in Michigan across wild mammal species.

| Species | Negative* | Positive* | Total* | % Positive full data | % Positive IHC-tested |
| --- | --- | --- | --- | --- | --- |
| Raccoon | 397 (383) | 510 (458) | 907 (841) | 56.2 | 54.5 |
| Striped skunk | 120 (108) | 73 (72) | 193 (180) | 37.8 | 40.0 |
| Gray fox | 32 (27) | 119 (115) | 151 (142) | 78.8 | 81.0 |
| Red fox | 111 (97) | 9 (9) | 120 (106) | 7.5 | 8.5 |
| Coyote | 43 (40) | 29 (29) | 72 (69) | 40.3 | 42.0 |
| American mink | 7 (5) | 11 (11) | 18 (16) | 61.1 | 68.8 |
| Gray wolf | 11 (11) | 4 (4) | 15 (15) | 26.7 | 26.7 |
| Black bear | 5 (5) | 1 (1) | 6 (6) | 16.7 | 16.7 |
| River otter | 3 (3) | 1 (1) | 4 (4) | 25.0 | 25.0 |
| Virginia opossum | 3 (3) | 0 (0) | 3 (3) | 0.0 | 0.0 |
| Bobcat | 2 (2) | 0 (0) | 2 (2) | 0.0 | 0.0 |
| Long-tailed weasel | 0 (0) | 2 (2) | 2 (2) | 100.0 | 100.0 |
| Silver fox | 2 (2) | 0 (0) | 2 (2) | 0.0 | 0.0 |
| American badger | 0 (0) | 1 (1) | 1 (1) | 100.0 | 100.0 |
| Fisher | 1 (1) | 0 (0) | 1 (1) | 0.0 | 0.0 |
| Short-tailed weasel | 0 (0) | 1 (1) | 1 (1) | 100.0 | 100.0 |
| Wolverine | 1 (1) | 0 (0) | 1 (1) | 0 | 0.00 |

\* number in parenthesis shows the number of IHC-tested cases

**Figure 2.**
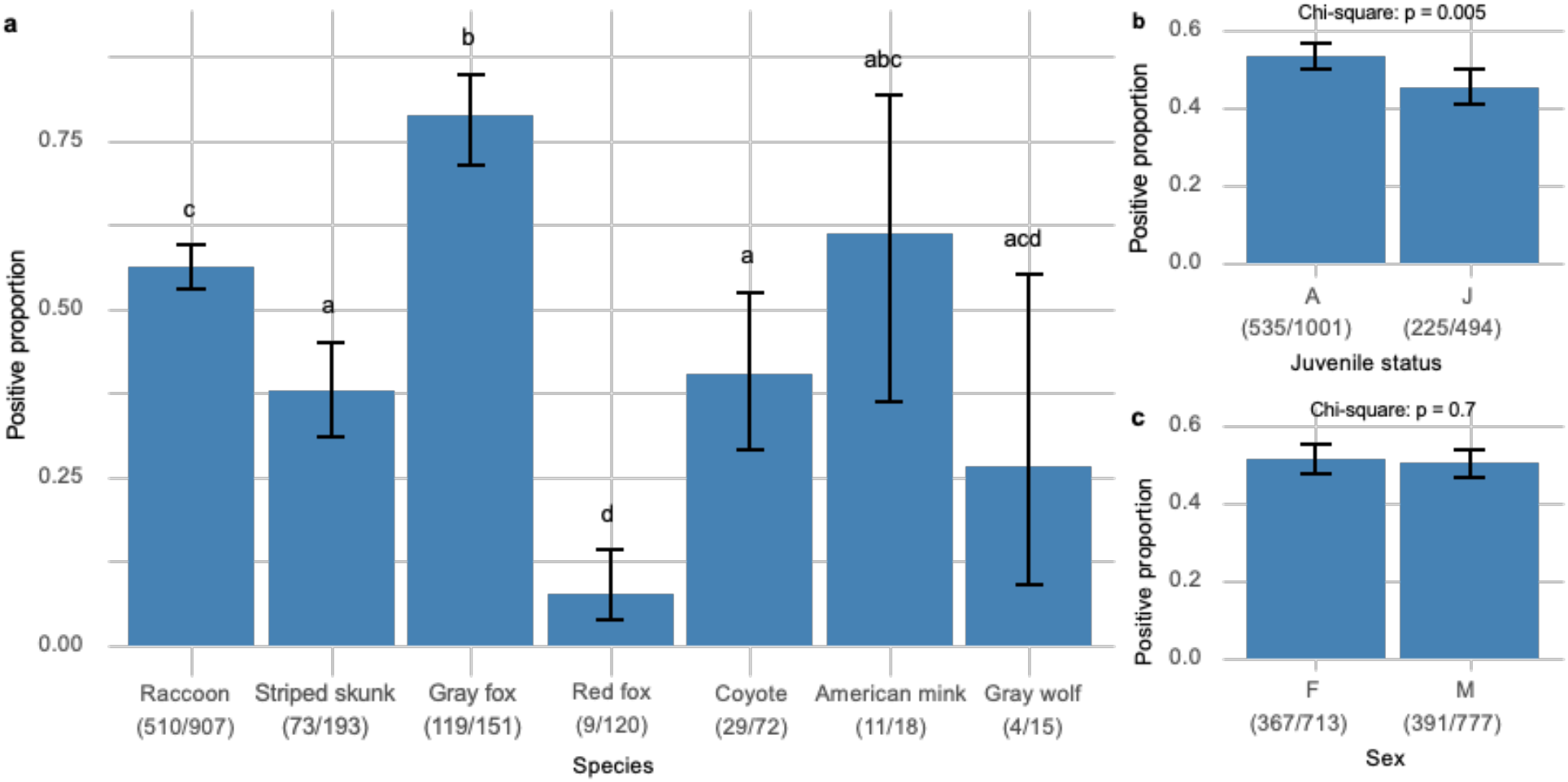
Positivity of CDV cases by host-associated category. (a) CDV positivity (proportion of positive submissions) across wildlife species with ≥ 10 submissions. (b) CDV positivity in adult (A) versus juvenile (J, < 1 year old) animals. (c) CDV positivity by sex. Positivity was calculated as the number of CDV-positive (confirmed or probable) submissions divided by the total number of submissions in each category. Error bars indicating 95% confidence intervals for the proportion. For panel (a), shared letters denote species that did not differ significantly in CDV positivity based on pairwise tests for equality of proportions. For panels (b) and (c), differences in CDV positivity between groups were assessed using chi-squared tests for equality of proportions.

Juvenile status was significantly associated with CDV test results (X^2^ = 7.9, p = 0.005) with juveniles having a positivity rate of 45.6%, lower than the 53.5% rate of adults (Fig 2b). In contrast, sex was not associated with test results (X^2^ = 0.15, p = 0.7); males had 51.5% while female had 50.3% positivity rates (Fig 2c).

Submissions from the Lower Peninsula accounted for the majority of cases (n = 1344), compared to the Upper Peninsula (n = 142). As described in Fitzgerald et al. [26], the number of submissions from the Upper Peninsula had a peak of more than 30 cases submitted in 2016 and 2017, and afterward, the number has remained at less than 10 cases each year (Fig S3). Positivity did not differ significantly between the two peninsulas: Lower Peninsula (50.8%) vs. Upper Peninsula (52.1%) (X^2^ = 0.04, p = 0.8). By county, the highest numbers of submissions were from Wayne County (n = 226) followed by Ingham County (n = 98). Positive animals were detected in 82 counties. The Bayesian CAR model estimated the mean county positivity to be 52.0% (95% credible interval: 38.4%–65.5%) (Fig 3). County-level posterior estimates ranged from 40.5% in St. Clair (95% CrI: 24.5%–56.1%) to 60.6% in Baraga (95% CrI: 43.4%–77.5%).

**Figure 3.**
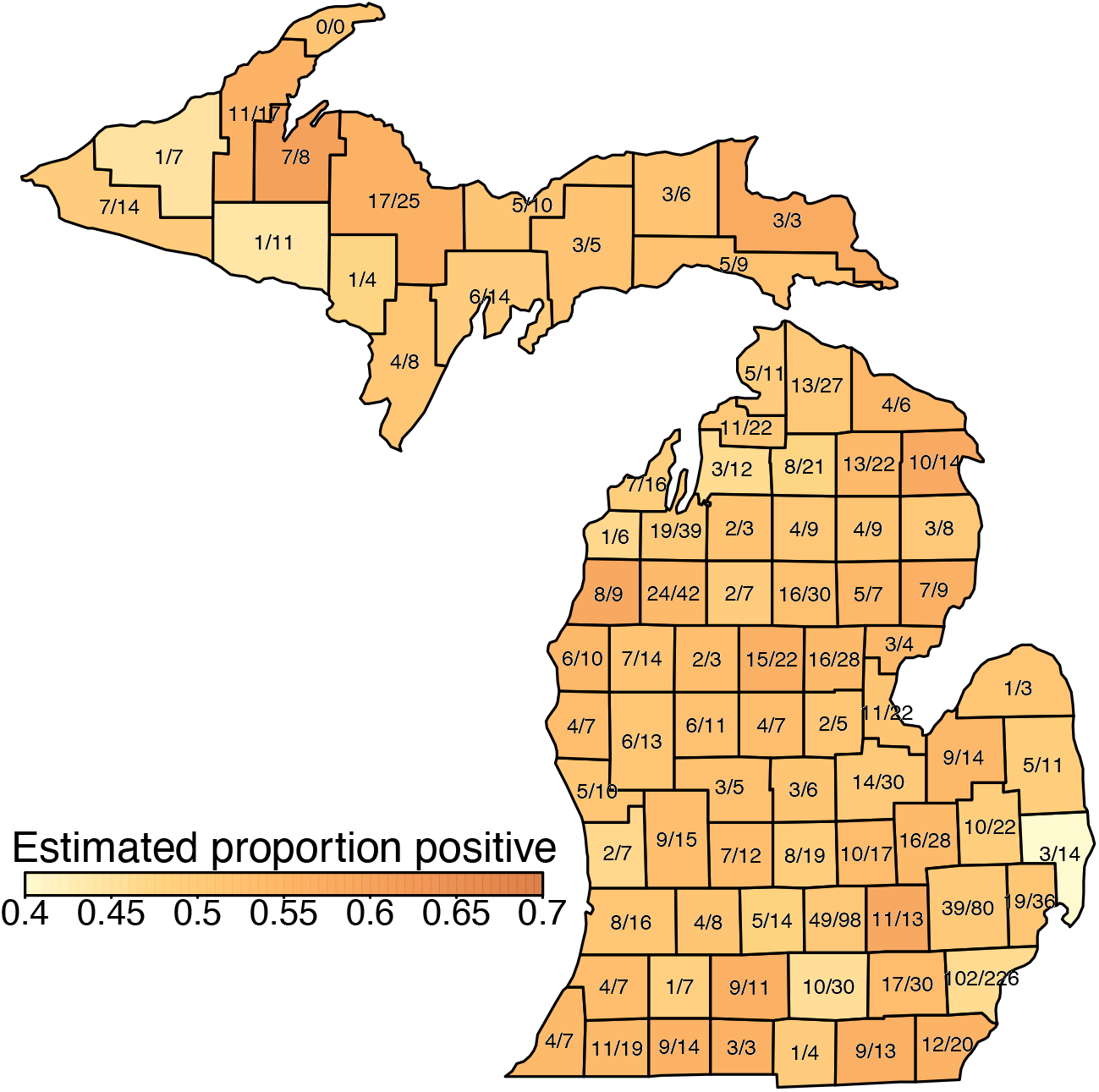
Geographic summary of CDV surveillance data. County-level map showing estimated CDV positivity per county calculated using a Bayesian conditional autoregressive model which accounts for variable sampling effort across counties and spatial correlation between neighboring counties. The total number of submissions and the number of CDV-positive samples are labeled for each county.

These estimates have large overlaps in their credible intervals suggesting that this data provides little evidence for large county-to-county differences in positivity.We next used a logistic regression to evaluate whether species, age, sex, peninsula, quarterly season, and year were significant predictors of CDV test results when combined. Reference categories for categorical variables were set as raccoon (species) and Q4 (Oct-Dec; quarter season). In the full model, quarterly season, species, juvenile status and year (included as a continuous variable) were significant predictors of CDV test results (Fig 4; Table S2). Compared to the reference category of racoons, gray foxes had higher odds for testing positive (odds ratio = 2.7, 95% CI: 1.8-4.2), while striped skunks, red foxes, gray wolfs, coyotes, and other species combined had lower odds. A general decrease in positivity over time was predicted with an estimated 0.97 annual fold change in odds ratio. Compared to reference category of Q4 (Oct-Dec), Q1 (Jan-Mar) and Q3 (Jul-Sep) had lower odds ratios.

**Figure 4.**
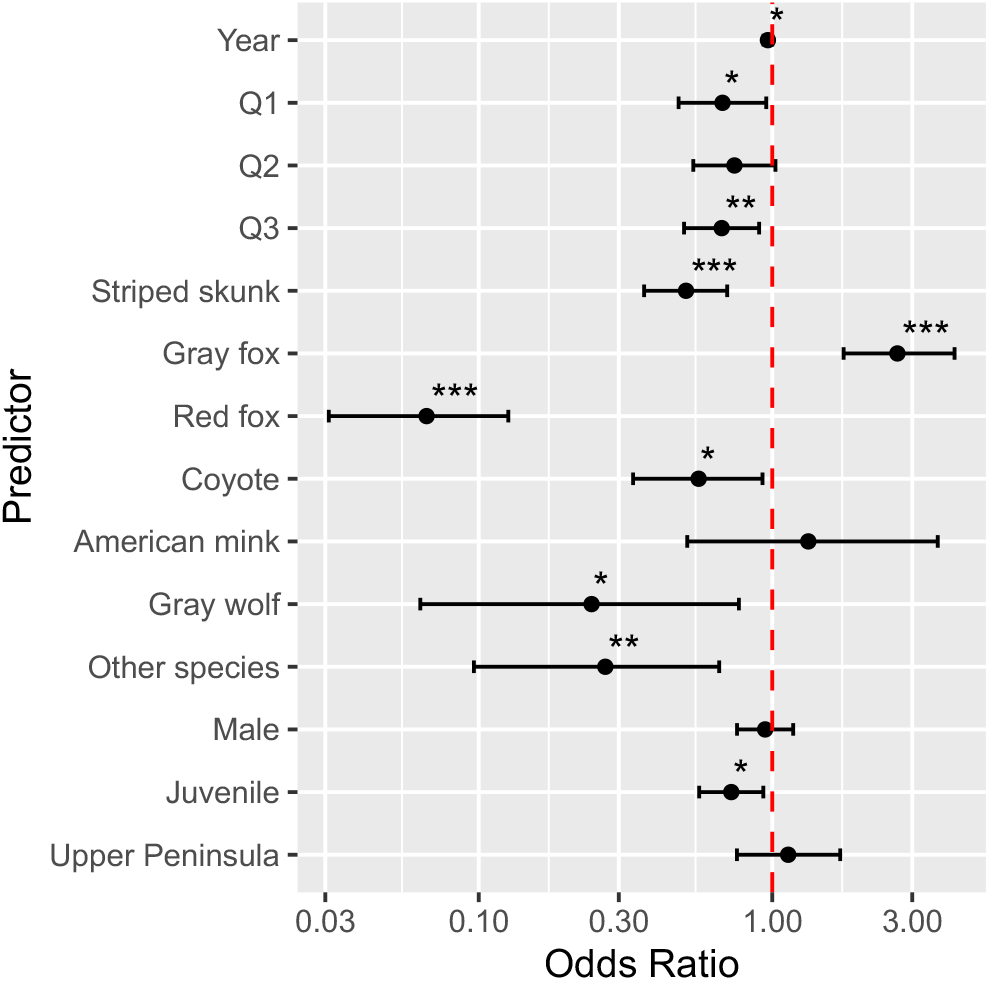
Predictors of CDV test positivity in Michigan wildlife based on a multivariable generalized linear model. Odds ratios (ORs) and 95% confidence intervals from a logistic regression evaluating the effects of species, juvenile status, sex, peninsula, quarterly season, and year on CDV test result (positive vs negative). Reference categories were raccoon for species and fourth quarter (Q4; Oct–Dec) for season. Points represent odds ratios and horizontal lines indicate 95% confidence intervals; OR > 1 indicates higher odds of testing positive relative to the reference category. Dashed red lines indicates an odds ratio of 1 equivalent to no difference and predictors significantly different from 1 are indicated with asterisk (*P* < 0.05, *; *P* < 0.01, **; *P* < 0.001, ***).

### Michigan wildlife and domestic dog sequences

This study generated 35 sequences of CDV from Michigan from both wild mesocarnivore species and domestic dogs. This includes 25 partial H gene sequences, 6 full-length H gene sequences, and 4 partial genomes (Table S3).

For wildlife samples, a total of 23 samples collected in 2022 and 2023 were sequenced for a partial H gene fragment (∼1100 bp). Sampled species included 13 racoons, four gray foxes, two minks, two skunks, one coyote, and one red fox. Four samples were further sequenced with ONT MinION to obtain longer contigs. De-novo assembly recovered contigs spanning most of the CDV genome for each sample; “23355_Skunk_2023” had the highest average read coverage at 205×, while the other samples had average coverages of 11-23× (Fig S5). Portions of the genome with low coverage and poor resolution were trimmed from the contigs. The accuracy of the assembled contigs was validated using a ∼1100 bp fragment of the H gene obtained by Sanger sequencing. The ONT-generated contigs matched the Sanger sequencing results exactly, demonstrating the accuracy of CDV sequences recovered from the ONT assemblies. One exception was observed at a single position in the “23040_Raccoon_2023” sample, where the Sanger sequence showed an ambiguous base R (A/G), but the ONT contig resolved it to adenine.

In addition to these 23 samples, one raccoon submitted to MSU VDL by a private veterinarian (i.e., not part of the MI DNR surveillance program) was sequenced, and three wildlife samples from 2019, originally reported by Fitzgerald et al. [26], were re-sequenced to obtain full-length H-gene sequences.

Eight domestic dog samples submitted to MSU VDL from 2021 to 2024 for CDV testing were also sequenced to investigate circulation between domestic dogs and wildlife animals. We recovered three full-length H genes and five partial H genes.

### CDV evolution across time and continents visualized by Nextstrain

To investigate local and global circulation of Michigan CDV lineages, we built a Nextstrain workflow. Our workflow consists of four datasets: a full-length H gene alignment and partial H gene alignment with or without recombinant sequences. Since no recombinant sequences were detected from Michigan, only recombinant free datasets are presented in the result section. The full length H gene dataset without potential recombinants contains 1,187 sequences including 10 new Michigan sequences, and the partial H gene dataset without potential recombinants contains 1,667 sequences including 35 new Michigan sequences.

Phylogenetic trees inferred from both the full-length H gene dataset and the partial H gene dataset showed largely concordant topologies (Fig S6). Both datasets classified each sequence into previously defined geographic lineages with high ultrafast bootstrap supports of ≥95 [48], resulting in the identification of 21 defined clades reported previously [11, 14, 16, 17, 24, 49–54] (Fig 5).

**Figure 5.**
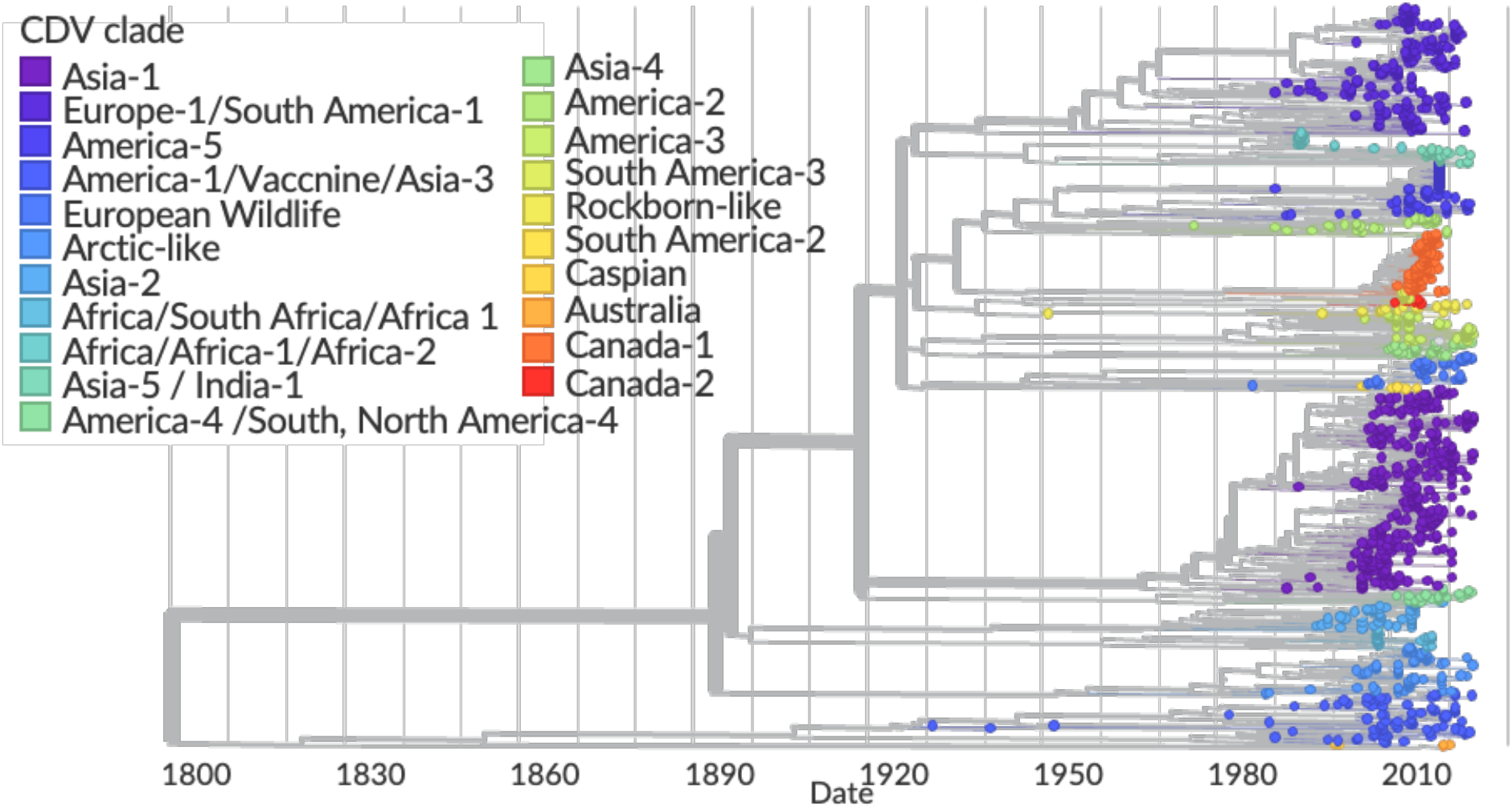
Time-scaled Nextstrain phylogeny of canine distemper virus (CDV) based on partial hemagglutinin (H) gene sequences. Time-resolved CDV phylogeny generated with Nextstrain using partial H gene sequences. Branch lengths are scaled to calendar time, showing the temporal distribution of lineages over the study period. Tree tips are color-coded by previously described geographic clades. An interactive version of this phylogeny, including full metadata and filtering options, is available on Nextstrain (https://nextstrain.org/community/nakakotanaka/nextstrain-cdv).

In both analyses, the America-1/Vaccine/Asia-3 clade, together with the Australian and Caspian lineages, consistently formed the most basal lineage. All remaining clades were contained within a single monophyletic group, with a time to most recent common ancestor (tMRCA) estimated at 1884 (95% CI: 1870–1917) in the full-length H gene dataset and 1893 (95% CI: 1877–1933) in the partial H gene dataset. Within this large monophyletic group, 14 clades clustered together as a well-supported sub-clade, whereas the remaining three clades (Africa/South Africa/Africa-1, Arctic-like, and Asia-2) were recovered either as poorly supported sub-clades (H gene dataset) or as distinct monophyletic groups (partial H gene dataset). These results suggest that the three latter clades are sister to the larger sub-clade consisting of 14 clades. Relationships across the 14 clades were less clearly resolved, frequently represented by polytomies or nodes with low bootstrap support.

In general, nodes in the partial H gene phylogeny exhibited lower bootstrap support, likely due to the reduced number of informative sites compared to the full-length H gene. This indicates that the full-length H gene likely provides more robust evolutionary relationships. At the same time, the partial H gene dataset adds complementary value by incorporating a larger number of sequences. This increased sampling allowed recognition of certain clades as monophyletic and independent clades rather than being a part of existing clades, such as Canada-1 and Canada-2, highlighting the utility of combining both datasets to obtain a more comprehensive understanding of CDV diversity (Fig 5).

### Three lineages of CDV are circulating in Michigan

In the context of the global CDV phylogeny, Michigan sequences were distributed across three clades: America-3, America-5, and Canada-1 (Fig 6). The America-3 clade includes 22 of the 35 Michigan sequences identified in this study, making it the most widespread clade in the state (Fig 6a). This lineage is composed primarily of U.S. sequences but also includes one sequence from a Mexican dog (KT266736.1) and one from a Canadian dog (OK666918.1). The U.S. sequences originate from Colorado, Texas, California, Missouri, and Wyoming, as well as from samples with unknown geographic information (Fig 6a). The estimated tMRCA of this lineage is 1976 (95% CI: 1968-1982) in the full-length H gene dataset and 1981 (95% CI: 1972-1995) in the partial H gene dataset. Within the America-3 clade, CDV sequences from 16 wild animals and four domestic dogs in Michigan formed a distinct sub-clade, suggesting localized circulation of this lineage. Within this Michigan subclade, dog and wildlife sequences were closely interspersed rather than forming host-specific clusters, indicating active transmission between domestic and wild hosts. tMRCA analysis indicates diversification of this Michigan sub-clade beginning in the early 2000s, with the full-length H gene dataset estimating a tMRCA of 2006 (95% CI: 2001-2010) and the partial H gene dataset estimating 2007 (95% CI: 2003-2012). The remaining two Michigan dog sequences in America-3 clustered separately: one formed a clade with a Canadian dog sequence imported from Aruba (OK666918), and the other formed a sub-clade with four U.S. dog sequences from Texas (MT932498.1, MT932496.1, MT932495.1, and MT932497.1). These cases indicate that some America-3 CDV lineages circulate across countries and states and may be distinct from the Michigan sub-clade locally circulating.

**Figure 6.**
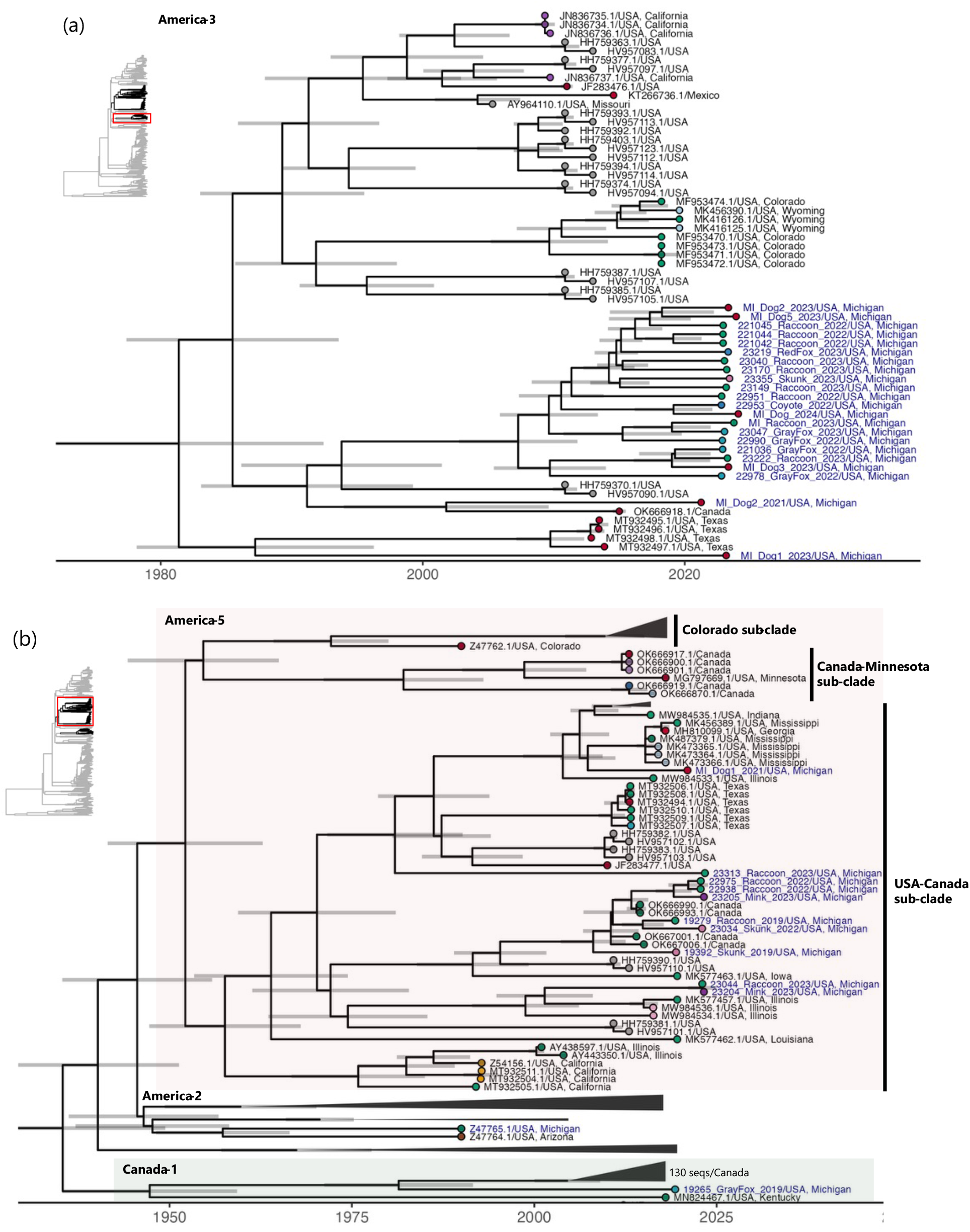
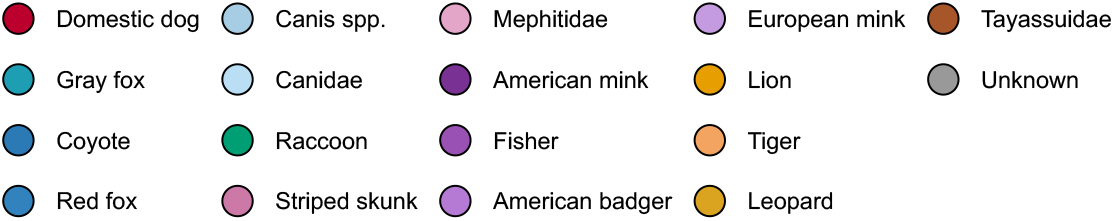
Clades containing Michigan sequences from a time-scaled Nextstrain phylogeny of canine distemper virus (CDV) based on partial hemagglutinin (H) gene sequences. (a) America-3 clade; (b) America-2, America-5, and Canada-1 clades. Branch lengths are scaled to calendar time, and gray bars at internal nodes represent 95% confidence intervals for node dates. Tree tips are color-coded by host species/taxonomic groups. Tip labels show accession number and country (state) of origin, with Michigan sequences highlighted in blue. Selected non-focal clades were collapsed to improve visualization of focal sequences. Inset at top left shows the full tree from Figure 5 indicating the region being shown here with a red box.

The America-5 clade includes 12 Michigan sequences from this study. Our analysis confirmed that the America-5 clade is a sub-clade of America-2 clade, as shown in previous studies [50, 55] (Fig 6b). America-5 mainly consists of sequences from USA (n = 88 in H gene/109 in partial H gene datasets) and 9 sequences from Canada in partial H gene datasets (Fig 6b). Within the America-5 clade, all 10 Michigan sequences fell within a group including Michigan sequences and sequences from multiple U.S. states and Canada (U.S.-Canada sub-clade) distinct from other American-5 subclades. Michigan sequences did not form monophyletic group within this U.S.-Canada sub-clade, indicating multiple introduction and active circulation of this lineage across U.S. states and Canada. Six Michigan wildlife samples were most closely related to four sequences from Canadian wild animals. As those Canadian sequences were from Ontario, a province geographically connected to Michigan, this sub-clade suggests movements of CDV and wildlife across the Michigan-Ontario border. Similar geographical clustering is observed between states. Michigan samples from raccoon (23044_Raccoon_2023) and mink (23204_Mink_2023) were most closely related to raccoon and skunk sequences from Illinois (MK577457.1, MW984536.1, MW984534.1). One Michigan raccoon sequence (23313_Raccoon_2023) was inferred to be a basal lineage of a sub-clade containing CDV in other states, such as Texas, Indiana, and California. One Michigan dog sequence lies within this multi-state clade and is most closely related to sequences from California and Georgia.

The Canada-1 clade was first reported by Giacinti et al., [50]. This clade originally consisted of Canadian sequences only, but our phylogeny suggests inclusion of two USA sequences to this clade: one sequence from a Kentucky raccoon (MN824467.1) and one sequence from a Michigan gray fox collected in 2019 (“19265_GrayFox_2019”) (Fig 6b). The Canada-1 clade is a sister of the America-2/America-5 clade. Fitzgerald et al. [26] previously reported this Michigan sequence as a “unique sequence type”, but subsequently reported genetic sequencing now allows placement of this sequence as a part of Canada-1 clade. This clade mainly comprises sequences from wildlife and is prevalent across Ontario. Notably, two sequences from the USA demonstrate a clear phylogenetic divergence from the Canadian sequences: a Kentucky sequence (MN824467.1) was placed as the most basal lineage in the Canada-1 clade, and the Michigan sequence was placed as a basal lineage of the Canadian sequences.

### America-3 clade has lower CT values than America-5 clade

We examined whether sample characteristics and phylogenetic clades explained variation in measured Ct values among wildlife and diagnostic samples (n = 31; Table S3). The full linear model included species, sex, age, clade, and specimen type as predictor variables. Geographic attributes were not included due to low sample number in each category. Among these, only clade was a significant predictor of Ct values (Table S4). Samples belonging to clade America-3 exhibited significantly lower Ct (i.e. higher estimated viral RNA abundance) values compared to clade America-5 (linear regression: mean difference = 7.5, 95% CI: 3.2-11.9, p = 0.002) (Fig 7).

**Figure 7.**
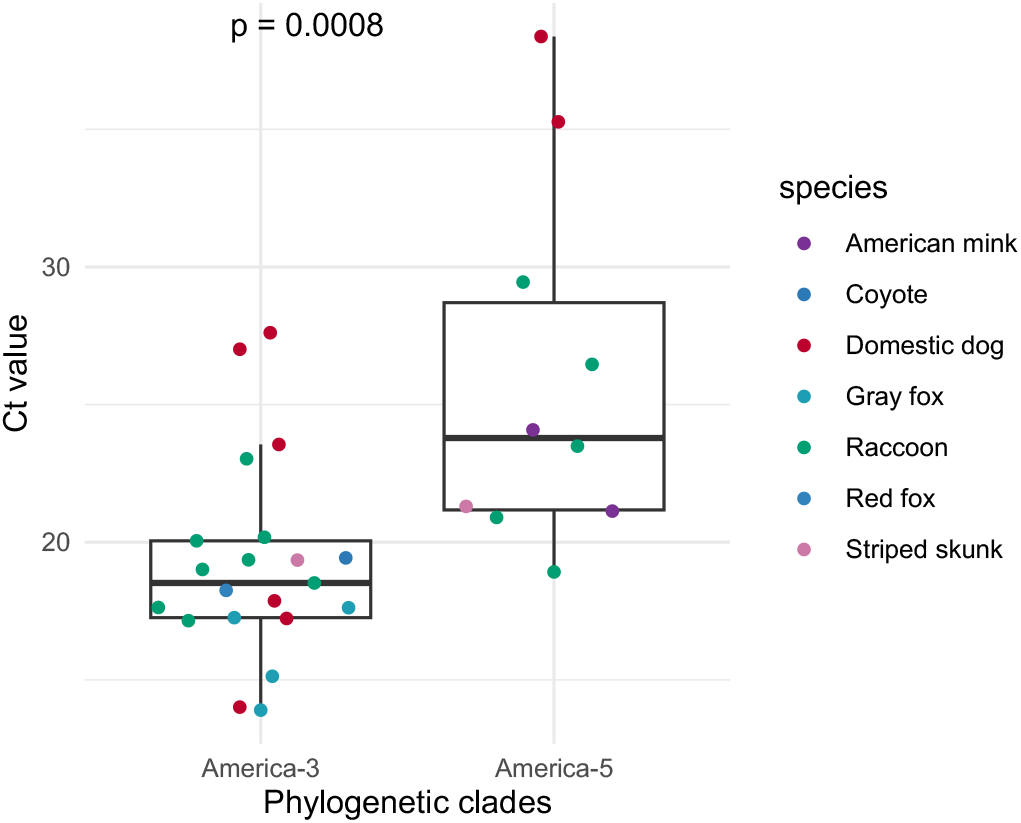
Ct value differences between America-3 and America-5 clades from Michigan CDV samples. Distribution of qRT-PCR Ct values for samples assigned to the America-3 and America-5 clades, with colors indicating host species. Differences in Ct values between clades were assessed using a Wilcoxon rank-sum test.

## Discussion

Our surveillance framework and phylogenetic analyses provide an integrated view of CDV circulation in Michigan over the past sixteen years. The data indicate that CDV is not an episodic problem in Michigan; rather, it is a persistent, spatially widespread pathogen maintained in common mesocarnivores and also detected in domestic dog populations. We identified temporal, geographic, and host-associated factors associated with CDV positivity in wild mammals and generated 35 new sequences from multiple wildlife species and domestic dogs.

Phylogenetic analysis revealed that three CDV lineages, America-3, America-5, and Canada-1, are currently circulating in Michigan. Differences in Ct values between America-3 and America-5 further suggest potential biological differences between these lineages and underscore the need for more detailed characterization in future studies. These findings highlight the importance of coupling wildlife surveillance with genomic sequencing to better detect lineage turnover and identify the emergence of new or epidemiologically important CDV lineages in a local context.

Species-level patterns in our dataset are consistent with CDV ecology described in other systems and provide a pragmatic basis for prioritizing wildlife health efforts in Michigan. In comparable wildlife surveillance systems in the United States and Canada, raccoons are often considered an important reservoir host, and large numbers of CDV-positive cases are frequently reported [50, 56]. Similarly, in our surveillance data, raccoons accounted for the largest number of submissions and CDV-positive cases. Canid hosts including gray foxes, coyotes, and red foxes are also commonly infected with CDV, and gray foxes in particular are reported to have high detection rate of CDV [57]. Consistent with this, we observed a higher positivity proportion in gray foxes and detected CDV-positive cases in coyotes and red foxes throughout the study period. Sustained CDV circulation in common mesocarnivores creates ongoing spillover risk into less frequently sampled or more vulnerable taxa. This risk is particularly relevant in Michigan given the presence of felid species of conservation concern that have been affected by CDV elsewhere, including cougar (*Puma concolor*) and Canada lynx (*Lynx canadensis*) [58, 59]. In practice, continued surveillance can provide an early warning system to trigger rapid investigation and response when unusual morbidity or mortality is detected in these species.

Temporal signals in our study further support the value of sustained surveillance. In wildlife systems, CDV dynamics can be episodic, with outbreaks recurring at multi-year intervals rather than remaining constant through time [60, 61]. Across our 16-year dataset, we observed an increase in CDV-positive detections during 2015–2018 in parallel with increased overall submissions [26], followed by lower but continued detection in in subsequent years. This pattern could reflect several non-mutually exclusive processes, including changes in reporting or submission effort, host demography and density [62], fluctuation in population immunity as susceptible individuals are replenished [63], viral lineage turnover [64].

We note that these data are complicated by the non-random sampling of animals. The positivity reported here are among animals reported to the MI DNR, due to displaying abnormal signs. To be reported an animal must interact with humans biasing results to animals found in close proximity to humans. Total reports will also vary with the public’s propensity to report abnormal behavior and spend time in areas containing wildlife. For example, counties with a high population will have more humans available for animal-human interaction but often less diverse/abundant wildlife. But these numbers do represent general trends in CDV observations and provide a measure of the presence and spread of this virus critical to animal health. We also note that our sequencing was limited to 2019–2023, which constrains inference about whether the increase in CDV detections around 2015 was accompanied by changes in circulating lineages. Nonetheless, these results provide a baseline for evaluating future outbreaks and for integrating expanded sequencing into routine surveillance to better inform management and response planning.

Sequencing and phylogenetic analysis support three epidemiologically relevant features of CDV in Michigan: (a) multiple co-circulating lineages across wild and domestic hosts rather than a single homogeneous enzootic lineage, (b) evidence of CDV exchange between wildlife and domestic dogs, and (c) connectivity with neighboring regions. Collectively, these features align with reports from other regions: CDV is a multi-host pathogen with frequent cross-species transmission, and its genetic diversity is usually geographically structured (often at continental scales) consistent with predominantly regional spread [14]. Periodic cross-border introductions can nonetheless seed new local transmission chains, supporting co-circulation of multiple lineages [16].

Phylogenetic intermixing of CDV sequences from domestic dogs and wildlife in Michigan suggest ongoing exchange at the dog–wildlife interface as observed in several previous studies. In the 1994 lion outbreak in the Serengeti in Tanzania, viruses found in lions were closely related to domestic dog strains suggesting spillover [7]. Long-term modeling further emphasizes frequent spillovers from dogs in this ecosystem [61]. In contrast, in many North American settings where routine vaccination in domestic dogs is more common, molecular investigations often frame wildlife as an important reservoir source for dog cases, with dog outbreaks tending to occur in under-vaccinated or high-risk populations, while wildlife circulation can persist independently [24, 65]. In Michigan, given generally high vaccination coverage in owned dogs, wildlife may represent an important source of exposure for domestic dogs, reinforcing the value of continued monitoring in wildlife populations.

We observed a clear signal of regional connectivity across neighboring U.S. states and Ontario, with Michigan sequences in all three detected lineages clustering with sequences from Canada and/or adjacent states. This pattern is consistent with predominantly regional spread of CDV facilitated by wildlife movement and human-mediated movement of domestic dogs. At the same time, the dominant lineages differed between jurisdictions: Canada-1 predominated in Ontario [50], whereas America-3 and America-5 predominated in Michigan, suggesting that cross-border links reflect intermittent introductions layered onto predominantly local transmission, rather than a single, well-mixed enzootic system spanning the border. Although the magnitude and direction of exchange cannot be inferred confidently without denser sampling, even intermittent cross-border movement can introduce new variants of concerns and shift local lineage composition, and co-circulation can create opportunities for homologous recombination [66]. Improving inference and situational awareness will therefore require coordinated sequencing and data sharing and analysis across state and provincial partners, particularly in border-adjacent areas and at interfaces where domestic dogs and wild carnivores are both represented.

Within Michigan, America-3 emerges as a lineage that warrants follow-up because of its high frequency in the sequenced dataset, evidence of local clustering, and molecular features that generate testable hypotheses. America-3 comprised 62.9% (22/35) of sequenced samples, and most grouped within a largely Michigan-only subclade, suggesting sustained local circulation during the years sampled. Among qRT-PCR–tested specimens, America-3 also exhibited lower Ct values than America-5, which could indicate lineage associated viral load differences. The Michigan America-3 subclade also carries several amino-acid substitutions of potential interest (F243S, I522V, Y549H; Fig S7). Y549H occurs at a residue implicated in SLAM (CD150) receptor interaction and has been linked in prior work to host association and/or virulence-related phenotypes in morbilliviruses, making it a plausible candidate for functional follow-up [53, 67]. I522V lies within the C-terminal globular head of H in a region implicated to associate with nectin-4 receptor engagement [68], whereas the functional relevance of F243S is currently unclear. Although sequence observations alone do not demonstrate altered transmissibility or virulence, the convergence of frequency, clustering, and receptor-proximal variation supports continued monitoring of America-3 and motivates focused phenotypic testing of this Michigan subclade.

Beyond the phylogenetic findings from Michigan, this study contributes an operational template for CDV genomic surveillance by implementing a reproducible Nextstrain workflow that can be updated as additional sequences and metadata accumulate. As sequencing becomes more accessible, the limiting step is increasingly not data generation but standardized analysis and interoperable metadata; a shared Nextstrain framework can improve temporal and geographic metadata consistency and make results more interpretable and comparable across laboratories and publications [25]. At present, our pipeline is implemented for hemagglutinin gene data, but extending the workflow to whole genomes and other genes is a high-priority next step because genome-scale data provide substantially greater phylogenetic resolution and stronger inference about introductions, persistence, and lineage turnover. Given that whole-genome CDV sequencing is already feasible using both short-read and nanopore platforms [51, 69], integrating WGS into routine surveillance is an achievable near-term goal. In parallel, the CDV field would benefit from a standardized, community-driven lineage nomenclature, similar to WHO’s measles genotype/strain naming system [70], to reduce ad hoc labeling and improve comparability across regions and time. Together, these advances would move CDV genomics from descriptive local strain tracking toward actionable surveillance: earlier identification of emergent or expanding lineages, clearer evidence for cross-jurisdiction connectivity, and a more defensible basis for outbreak response, particularly when events threaten conservation-relevant wildlife or vulnerable domestic dog populations.

## Author contributions

Conceptualization: JM, SF, KD, RM, SSM, KN; Data curation: JM, SF, LH, KN, SA, KCH, AW, DT; Formal analysis: KN, SSM; Funding acquisition: KD, RM, JM; Investigation: JM, LH, KN, SA, KCH, AW, DT; Methodology: KN, JM, SF, LH, SA, KCH, AW; Project administration: KN, SSM, JM, LH, SA, KCH, KD, RM; Resources: JM, SF, LH, SA, KCH, AW, DT, TTN, KD, RM, KN, SSM; Software: KN, SSM, KCH; Supervision: KD, RM, SSM, TTN; Validation: KN, SSM, JM, LH, SA, KCH, AW, DT, TTN, KD, RM; Visualization: KN, SSM, Writing – Original draft: KN, SSM; Writing – review & editing: KN, SSM, JM, LH, SA, KCH, AW, DT, TTN, KD, RM, SF

## Acknowledgements

We thank the individuals and communities who reported animals exhibiting CDV-like signs to the MI DNR, and the MI DNR staff and field personnel who collected samples used in this study. We also thank virology section staff at MSU VDL for conducting RNA-extraction, qRT-PCR, and retention of samples for this study. This work was supported in part through computational resources and services provided by the Institute for Cyber-Enabled Research at Michigan State University.

## Ethical Statement

This study used retrospective diagnostic and surveillance submissions to the Michigan State University Veterinary Diagnostic Laboratory and the Michigan Department of Natural Resources Wildlife Disease Laboratory. No animals were prospectively enrolled or sampled for research purposes. Ethical review and approval were not required in accordance with institutional policy.

