## Supplementary document for "Epidemiology and phylodynamic analysis of canine distemper virus circulating in Michigan, USA"

**Title:**

**Affiliation:**

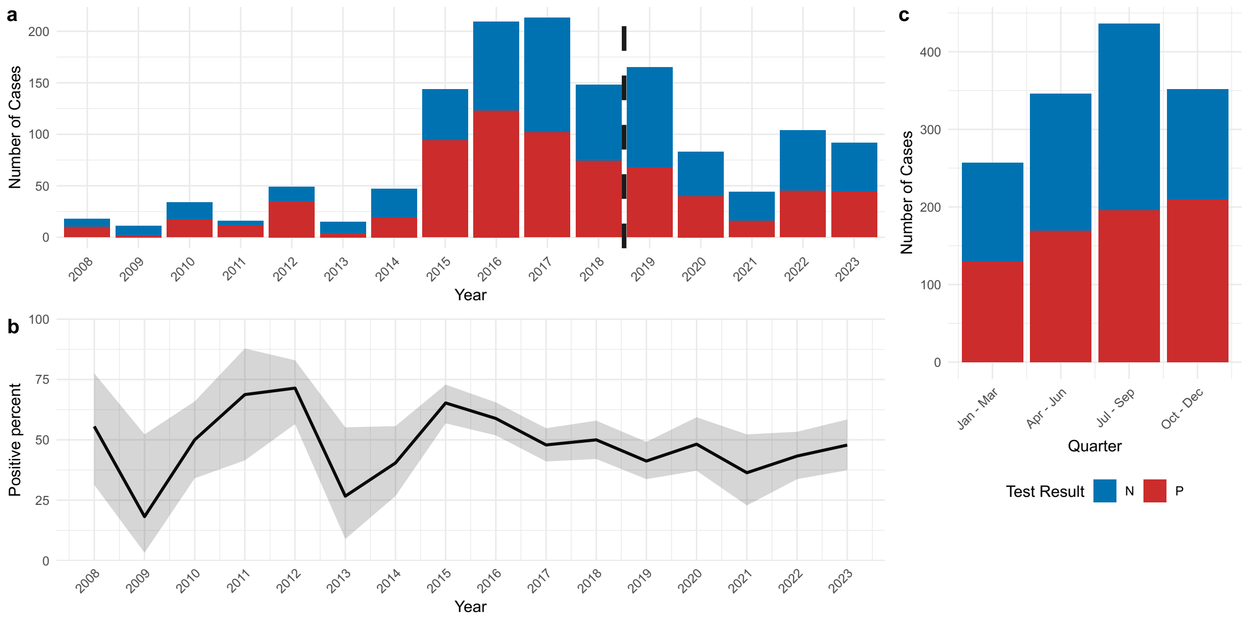


Figure S1. **Temporal dynamics of CDV surveillance data from 2008 to 2023 from only IHC-tested dataset.** (a) Annual numbers of wildlife submissions and immunohistochemistry (IHC) test results, showing counts of IHC-positive, IHC-negative cases. Dashed line indicates the break between cases previously reported in Fitzerald et al. 2022 and cases newly reported here. (b) Temporal changes in CDV positivity, expressed as the proportion of positive submissions among all tested samples with confidence interval. (c) Total number of submissions summarized by calendar quarter across the study period.

**Fitzgerald SD, Melotti JR, Cooley TM, Wise AG, Maes RK, *et al.*** Geographic spread of canine distemper in wild carnivores in michigan, usa: pathology and epidemiology, 2008–18. *J Wildl Dis* 2022;58:562–574.

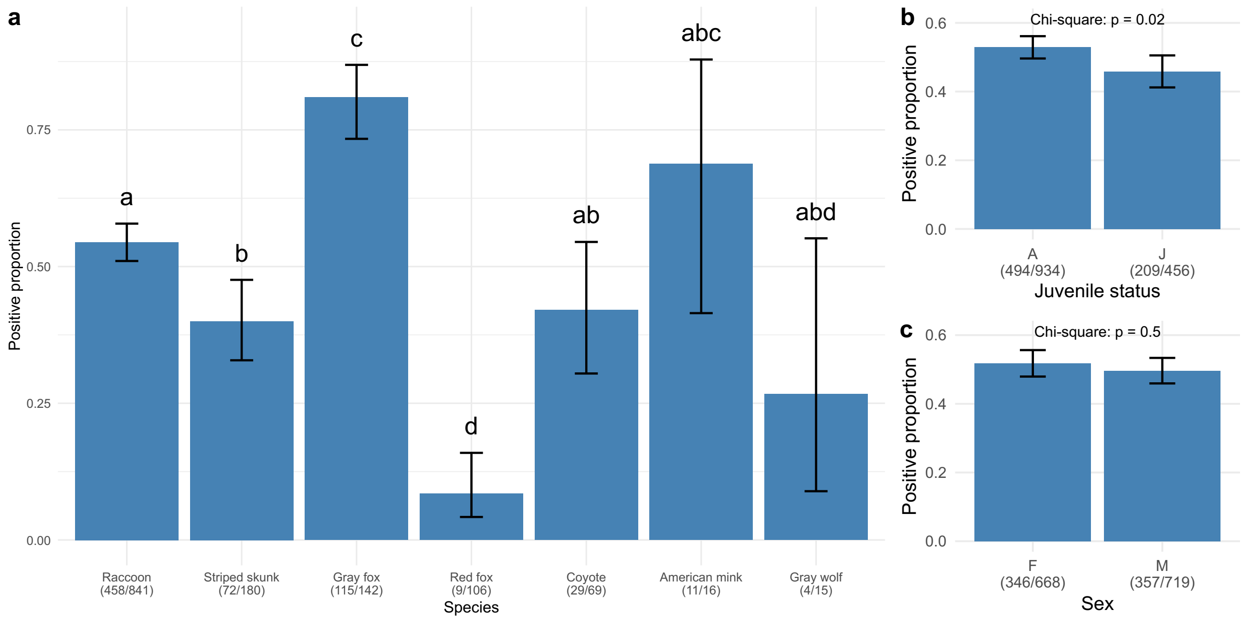


**Figure S2. Positivity of CDV cases by host-associated category from IHC-tested cases.** (a) CDV positivity (proportion of positive submissions) across wildlife species with ≥ 10 submissions. (b) CDV positivity in adult (A) versus juvenile (J, < 1 year old) animals. (c) CDV positivity by sex. Positivity was calculated as the number of IHC-tested CDV-positive submissions divided by the total number of IHC-tested submissions in each category. Error bars indicating 95% confidence intervals for the proportion). For panel (a), shared letters denote species that did not differ significantly in CDV positivity based on pairwise tests for equality of proportions. For panels (b) and (c), differences in CDV positivity between groups were assessed using chi-squared tests for equality of proportions.

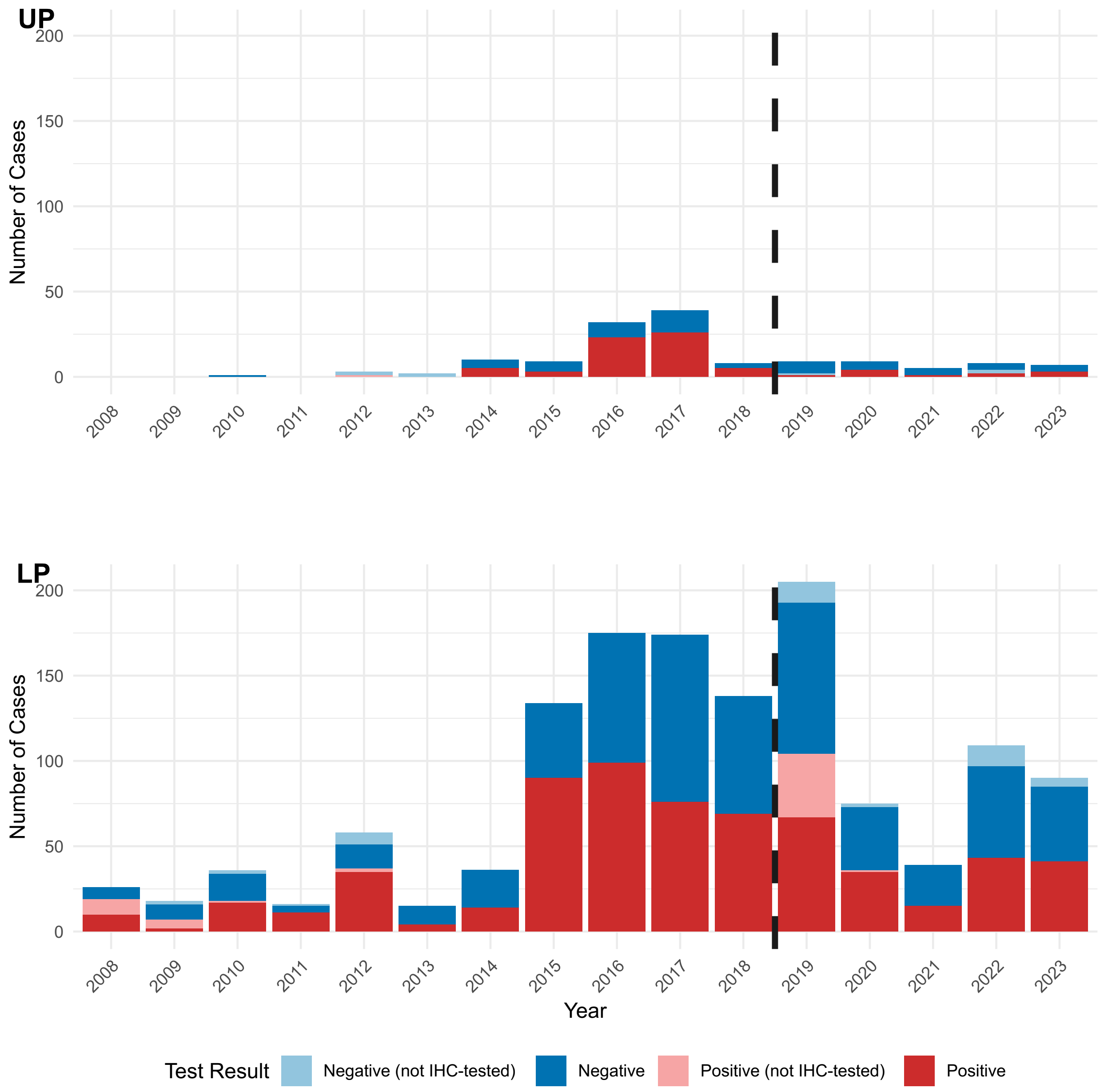


**Figure S3. Geographic summary of CDV surveillance data by Upper and Lower Peninsula.** Counts of submissions and CDV test results for counties (a) in the Upper Peninsula and (b) in the Lower Peninsula. Dashed line indicates the break between cases previously reported in Fitzerald et al. 2022 and cases newly reported here.

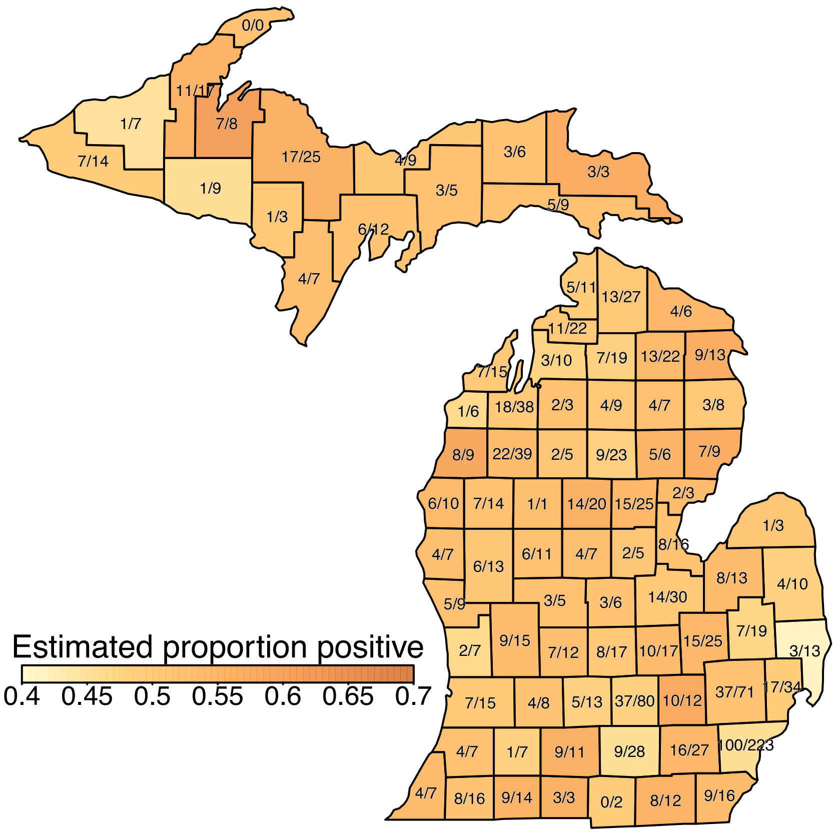


**Figure S4. Geographic summary of CDV surveillance using only IHC-tested data.** County-level map showing estimated CDV positivity (proportion of positive submissions) per county calculated using a Bayesian conditional autoregressive model which accounts for variable sampling effort across counties and spatial correlation between neighboring counties. The total number of submissions and the number of CDV-positive samples are labeled for each county.

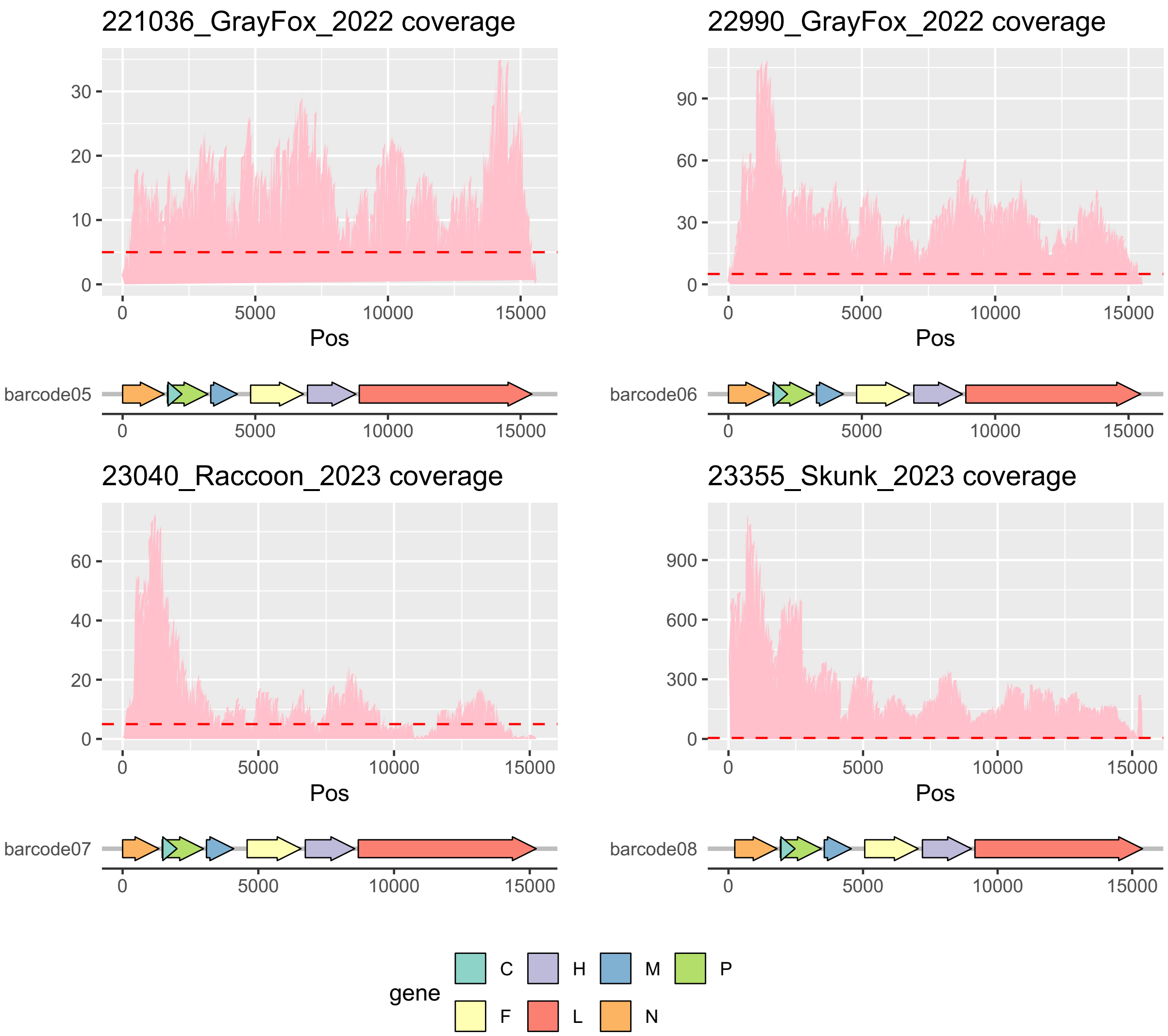


**Figure S5. Coverage profiles of Oxford Nanopore MinION generated CDV genomes from four samples.** Red line indicates 10x coverage.

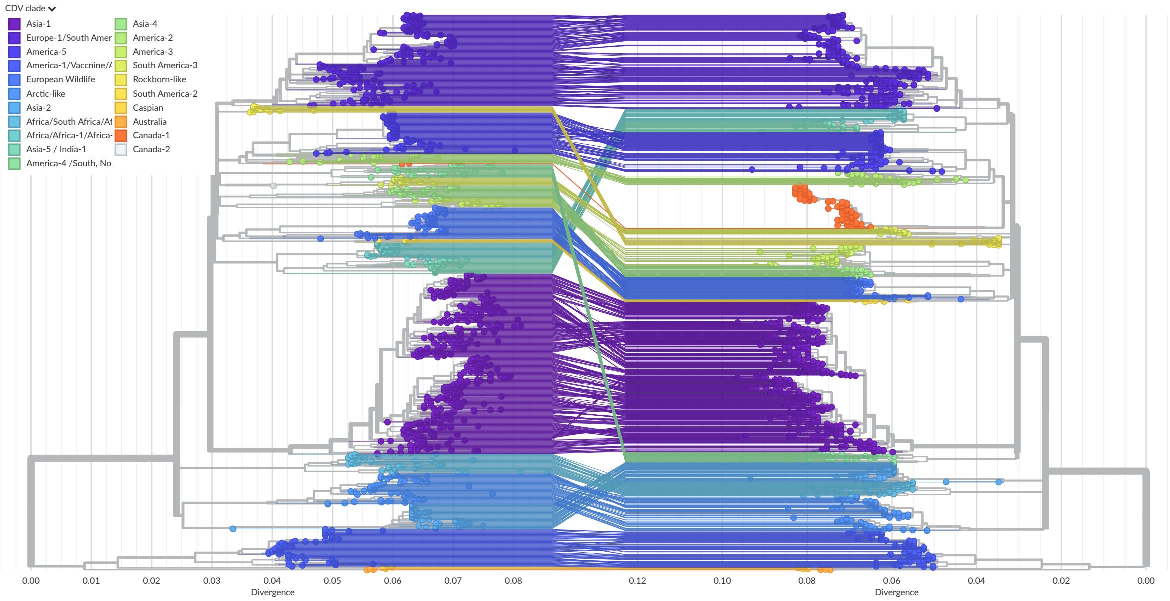


**Figure S6. Concordance between phylogenetic trees inferred from full-length and partial hemagglutinin (H) gene sequences of canine distemper virus (CDV).**

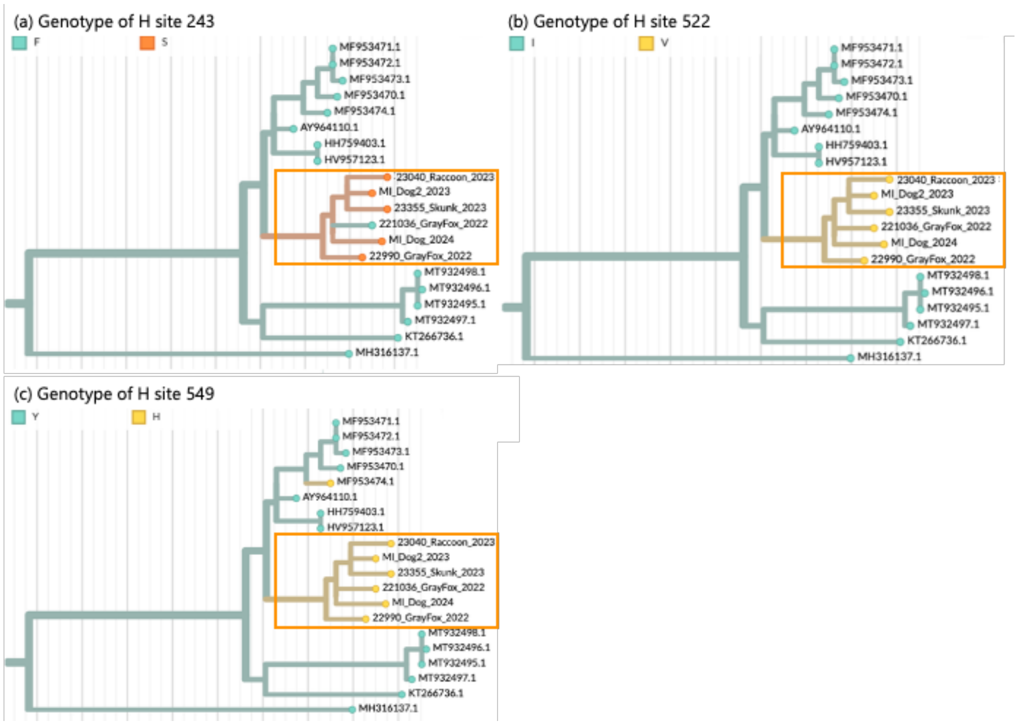


**Figure S7. Nextstrain visualization of homoplastic amino acid substitutions in the America-3 Michigan subclade.** The same America-3 canine distemper virus H gene phylogeny is shown in each panel, with tips colored by amino acid state at positions 243 (a), 522 (b), and 549 (c). These three substitutions, F243S, I522V, and Y549H, were identified as inferred homoplastic mutations when selecting the Michigan subclade node in Nextstrain. The Michigan subclade is indicated by orange squares.

**Table S1. Primers used for CDV H gene amplification and sequencing**

| Primer name | Direction | Sequence | Reference | Notes |
|---|---|---|---|---|
| CDVff1 | forward | TCGAAATCCTATGTGAGATCACT | Lan et al. 2006 | - |
| CDVHS2 | reverse | ATGCTGGAGATGGTTTAATTCAATCG | Lan et al. 2006 | - |
| CDVHS1 | forward | AACTTAGGGCTCAGGTAGTCC | Lan et al. 2006 | - |
| CDVHforD | forward | GACACTGGCTTCCTTGTGTGTAG | Lan et al. 2006 | - |
| CDVHr2 | reverse | GTTCTTCTTGTTTCTCAGAGG | Lan et al. 2006 | - |
| CDVP2F | forward | ACTTCCGCGATCTCCACT | Pardo et al.2005 | - |
| CDVP3R | reverse | ACACTCCGTCTGAGATAGC | Pardo et al.2005 | - |
| CDVP5R | reverse | GTGAACTGGTCTCCTCTA | Pardo et al.2005 | - |
| CDV H 381 F | forward | GAACAGGGAGTTCGACTTCC | - | designed in-house |
| CDV-1 Fwd 7480 | forward | CGATCTCCACTGGTGCATTA | - | designed in-house |
| CDV - 2 Fwd 7620 | forward | ACATATTCCCACCATACAGATGCA | - | designed in-house |
| CDV H 552 F | forward | ACCATACAGATGCAGTGGAG | - | designed in-house |
| CDV H 734R | reverse | GTGTCGAACTCCCCTTCAAT | - | designed in-house |
| CDV H 1364R | reverse | TTGGGAGGAATGGTAAGCC | - | designed in-house |
| CDV H 884F | forward | TGTGTGTAGATGAGAGCACC | - | designed in-house |
| CDV H 1068R | reverse | ATCTTTGATGAARCCACGGT | - | designed in-house |
| CDV - 2.2 Rev 8343 | reverse | TTCAGTATAACMGGACCGTATGTT | - | designed in-house |
| CDV H 1364R | reverse | TTGGGAGGAATGGTAAGCC | - | designed in-house |
| CDV H 1541 R | reverse | GGCAACACCACTAAATTGGA | - | designed in-house |
| 204+ | forward | GAATTCGACTTCCGCGATCTCC | Kapil et al. 2008 | Used in Fitzgerald et al. 2022 to amplify ~1160bp H gene |
| 232b- | reverse | TAGGCAACACCACTAATTTRGACTC | Kapil et al. 2008 | |

Lan NT, Yamaguchi R, Kawabata A, Uchida K, Kai K, Sugano S, Tateyama S. Stability of canine distemper virus (CDV) after 20 passages in Vero-DST cells expressing the receptor protein for CDV. Vet Microbiol. 2006 Dec 20;118(3-4):177-88. doi: 10.1016/j.vetmic.2006.07.015. Epub 2006 Sep 18. PMID: 16982161.

Pardo ID, Johnson GC, Kleiboeker SB. Phylogenetic characterization of canine distemper viruses detected in naturally infected dogs in North America. J Clin Microbiol. 2005 Oct;43(10):5009-17. doi: 10.1128/JCM.43.10.5009-5017.2005. PMID: 16207955; PMCID: PMC1248462.

Kapil S, Allison RW, Johnston L 3rd, Murray BL, Holland S, Meinkoth J, Johnson B. Canine distemper virus strains circulating among North American dogs. Clin Vaccine Immunol. 2008 Apr;15(4):707-12. doi: 10.1128/CVI.00005-08. Epub 2008 Feb 6. PMID: 18256210; PMCID: PMC2292659.

**Table S2. Multivariable generalized linear model of predictors of CDV test positivity in Michigan wildlife, 2008–2023 using full dataset.** Generalized linear model with binomial distribution and logit link; outcome variable is CDV test result (positive vs negative). Values shown are odds ratios (OR), 95% confidence intervals (CI), and p-values. Reference categories: raccoon (species), adult (juvenile status), female (sex), Lower Peninsula (peninsula), and Q4 (Oct–Dec) for season. OR > 1 indicates higher odds of testing CDV-positive relative to the reference category.

| Predictor | | Estimated odds ratio (log scale) | Standard error | Low CI | High CI | p-value |
|---|---|---|---|---|---|---|
| **(Intercept)** | | 2.72 | 0.23 | 1.73 | 4.31 | $2 \times 10^{-5}$ |
| Species | American mink | 1.33 | 0.49 | 0.51 | 3.66 | 0.6 |
| | **Coyote** | 0.56 | 0.26 | 0.34 | 0.93 | 0.03 |
| | **Gray fox** | 2.67 | 0.22 | 1.75 | 4.18 | $9 \times 10^{-6}$ |
| | **Gray wolf** | 0.24 | 0.62 | 0.06 | 0.77 | 0.02 |
| | **other** | 0.27 | 0.48 | 0.10 | 0.66 | 0.01 |
| | **Red fox** | 0.07 | 0.36 | 0.03 | 0.13 | $2 \times 10^{-14}$ |
| | **Striped skunk** | 0.51 | 0.17 | 0.37 | 0.70 | $5 \times 10^{-5}$ |
| Age | **Juvenile** | 0.73 | 0.13 | 0.56 | 0.93 | 0.01 |
| Sex | Male | 0.95 | 0.11 | 0.76 | 1.18 | 0.6 |
| Peninsula | Upper Peninsula | 1.13 | 0.21 | 0.76 | 1.70 | 0.5 |
| **Quarter** | **Q1** | 0.68 | 0.18 | 0.48 | 0.95 | 0.03 |
| | Q2 | 0.74 | 0.16 | 0.54 | 1.03 | 0.07 |
| | **Q3** | 0.67 | 0.15 | 0.50 | 0.90 | 0.01 |
| **Year** | | 0.97 | 0.02 | 0.93 | 1.00 | 0.05 |

**Table S3. Wildlife and diagnostic CDV samples from Michigan sequenced in this study.** For each sample, sample name, host species, specimen type, year, Michigan county, age, sex, sequencing platform, viral gene region(s) sequenced, and sequence length (bp) are shown. Age class: A, adult; J, juvenile (< 1 year); –, not recorded. Sex: M, male; F, female; –, not recorded. "Wildlife" and "Diagnostics" indicate sample origin categories. UP, Upper Peninsula; ONT, Oxford Nanopore Technologies; p, partial gene region (e.g., pH = partial hemagglutinin gene, pL = partial L gene). Where two sequence lengths are listed, the first corresponds to ONT data and the second to Sanger sequences. A dash in the county column indicates county of origin was not recorded. Two sequences with asterisks are either short or fragmented, thus not included in the phylogenetic work.

| Project | Sample Name | Species | Specimen Type | Year | Michigan County | Age | Sex | Sequence Method | Sequenced Region | Genbank Accession | Sequence Length | Lineage | RT-PCR Ct value |
|---|---|---|---|---|---|---|---|---|---|---|---|---|---|
| Wildlife | 19265_GrayFox_2019 | Gray Fox | Lung | 2019 | Ingham | A | M | Sanger | H | PX789675 | 1964 | Canada-1 | - |
| Wildlife | 19279_Raccoon_2019 | Raccoon | Lung | 2019 | Oakland | J | F | Sanger | H | PX789676 | 2003 | America-5 | - |
| Wildlife | 19392_Skunk_2019 | Skunk | Lung | 2019 | Wayne | A | F | Sanger | H | PX789677 | 2045 | America-5 | - |
| Wildlife | 22953_Coyote_2022 | Coyote | Lung | 2022 | Kalkaska | J | M | Sanger | pH | PX789686 | 1097 | America-3 | 19.43 |
| Wildlife | 221036_GrayFox_2022 | Gray Fox | Lung | 2022 | Clare | A | F | ONT, Sanger | pN, P, C, H, M, F, H, L | PX789704 | 15040, 1097 | America-3 | 13.9 |
| Wildlife | 22978_GrayFox_2022 | Gray Fox | Lung | 2022 | Houghton (UP) | A | F | Sanger | pH | PX789688 | 1076 | America-3 | 17.26 |
| Wildlife | 22990_GrayFox_2022 | Gray Fox | Lung | 2022 | Ionia | J | M | ONT, Sanger | pN, P, C, H, M, F, H, pL | PX789705 | 15284 , 1098 | America-3 | 15.13 |
| Wildlife | 22951_Raccoon_2022 | Raccoon | Lung | 2022 | Clare | J | F | Sanger | pH | PX789694 | 1098 | America-3 | 19.36 |
| Wildlife | 22975_Raccoon_2022 | Raccoon | Lung | 2022 | Genesee | A | M | Sanger | pH | PX789695 | 1111 | America-5 | 26.46 |
| Wildlife | 221045_Raccoon_2022 | Raccoon | Lung | 2022 | Ingham | A | F | Sanger | pH | PX789696 | 1117 | America-3 | 17.15 |
| Wildlife | 22938_Raccoon_2022 | Raccoon | Lung | 2022 | Lapeer | A | M | Sanger | pH | PX789698 | 1103 | America-5 | 23.49 |
| Wildlife | 221042_Raccoon_2022 | Raccoon | Lung | 2022 | Livingston | A | M | Sanger | pH | PX789699 | 1100 | America-3 | 20.05 |
| Wildlife | 23045_Raccoon_2022 | Raccoon | Lung | 2022 | Monroe | A | M | Sanger | pH* | PX789700 | 761 | America-5 | 29.45 |
| Wildlife | 221044_Raccoon_2022 | Raccoon | Lung | 2022 | Washtenaw | A | M | Sanger | pH | PX789701 | 1103 | America-3 | 20.18 |
| Wildlife | 23034_Skunk_2022 | Skunk | Lung | 2022 | Wayne | A | F | Sanger | pH | PX789702 | 1101 | America-5 | 21.3 |
| Wildlife | 23219_RedFox_2023 | Red Fox | Lung | 2023 | Livingston | J | M | Sanger | pH | PX789687 | 1101 | America-3 | 18.25 |
| Wildlife | 23047_GrayFox_2023 | Gray Fox | Lung | 2023 | Kent | A | M | Sanger | pH | PX789689 | 1103 | America-3 | 17.62 |
| Wildlife | 23205_Mink_2023 | Mink | Lung | 2023 | Lapeer | J | F | Sanger | pH | PX789690 | 1103 | America-5 | 24.08 |
| Wildlife | 23204_Mink_2023 | Mink | Lung | 2023 | Oakland | A | M | Sanger | pH | PX789691 | 1103 | America-5 | 21.13 |
| Wildlife | 23044_Raccoon_2023 | Raccoon | Lung | 2023 | Berrien | A | F | Sanger | pH | PX789692 | 1121 | America-5 | 20.9 |
| Wildlife | 23170_Raccoon_2023 | Raccoon | Lung | 2023 | Cass | A | M | Sanger | pH | PX789693 | 1101 | America-3 | 23.03 |
| Wildlife | 23149_Raccoon_2023 | Raccoon | Lung | 2023 | Jackson | A | M | Sanger | pH | PX789697 | 1098 | America-3 | 18.52 |
| Wildlife | 23040_Raccoon_2023 | Raccoon | Lung | 2023 | Wayne | A | F | ONT, Sanger | P, C, H, F, H, pL | PX789706 | 8079 , 1117 | America-3 | 17.63 |
| Wildlife | 23222_Raccoon_2023 | Raccoon | Lung | 2023 | Wexford | A | M | Sanger | pH | PX789673 | 1103 | America-3 | 19.01 |
| Wildlife | 23313_Raccoon_2023 | Raccoon | Lung | 2023 | Oceana | A | M | Sanger | pH | PX789703 | 1103 | America-5 | 18.92 |
| Wildlife | 23355_Skunk_2023 | Skunk | Lung | 2023 | Eaton | A | F | ONT, Sanger | N, P, C, H, M, F, H, pL | PX789707 | 15157 , 1063 | America-3 | 19.35 |
| Diagnostics | MI_Dog1_2021 | Dog | Serum | 2021 | - | J | M | Sanger | H | PX789678 | 1941 | America-5 | 35.27 |
| Diagnostics | MI_Dog2_2021 | Dog | Blood | 2021 | - | J | F | Sanger | pH | PX789679 | 1011 | America-3 | 27.61 |
| Diagnostics | MI_Dog1_2023 | Dog | Lung | 2023 | - | J | M | Sanger | pH | PX789680 | 1044 | America-3 | 27.01 |
| Diagnostics | MI_Dog2_2023 | Dog | Blood | 2023 | - | J | M | Sanger | H | PX789681 | 2008 | America-3 | 14.01 |
| Diagnostics | MI_Dog3_2023 | Dog | Serum | 2023 | - | J | M | Sanger | pH | PX789682 | 1034 | America-3 | 23.55 |
| Diagnostics | MI_Dog4_2023 | Dog | Blood | 2023 | - | J | F | Sanger | pH* | PX789674 | 649, 674 | America-5 | 38.37 |
| Diagnostics | MI_Raccoon_2023 | Raccoon | Lung | 2023 | - | - | - | Sanger | pH | PX789683 | 1046 | America-3 | 18.25 |
| Diagnostics | MI_Dog5_2023 | Dog | Lung | 2023 | - | A | M | Sanger | pH | PX789684 | 1067 | America-3 | 17.23 |
| Diagnostics | MI_Dog_2024 | Dog | Serum | 2024 | - | J | F | Sanger | H | PX789685 | 2006 | America-3 | 17.87 |

**Table S4. Multivariable linear regression of qRT-PCR Ct values in Michigan canine distemper virus (CDV) samples.** Linear regression model evaluating associations between qRT-PCR Ct values and host species, viral clade, sex, specimen type, and age. For categorical predictors, estimates represent the difference in Ct value relative to the reference categories: Coyote species, America-3 clade, female sex, adult age, and whole blood specimen type.

| Predictor | | Estimate | Standard error | p value |
|---|---|---|---|---|
| **(Intercept)** | | **17.53** | **7.63** | **0.03** |
| Species | Domestic Dog | 3.97 | 5.74 | 0.5 |
| | Gray Fox | -1.40 | 5.65 | 0.8 |
| | Mink | -2.96 | 6.20 | 0.6 |
| | Raccoon | 1.20 | 5.51 | 0.8 |
| | Red Fox | -1.18 | 6.42 | 0.9 |
| | Skunk | -0.06 | 6.75 | 1.0 |
| **Clade** | **America-5** | **7.54** | **2.07** | **0.002** |
| Sex | Male | 0.25 | 1.96 | 0.9 |
| Specimen | Lung | -0.91 | 4.59 | 0.8 |
| | Serum | -1.18 | 3.77 | 0.8 |
| Age | Juvenile | 2.57 | 2.84 | 0.4 |
